# Cardio-centric hemodynamic management improves spinal cord oxygenation and mitigates hemorrhage in acute spinal cord injury

**DOI:** 10.1101/2020.03.29.014498

**Authors:** Alexandra M. Williams, Neda Manouchehri, Erin Erskine, Keerit Tauh, Kitty So, Femke Streijger, Katelyn Shortt, Kyoung-Tae Kim, Brian K. Kwon, Christopher R. West

## Abstract

Chronic high-thoracic and cervical spinal cord injury (SCI) results in a complex phenotype of cardiovascular consequences, including impaired left-ventricular contractility. Here, we sought to determine whether such dysfunction manifests immediately post-injury, and if so, whether correcting impaired contractility can improve spinal cord oxygenation (SCO_2_), blood flow (SCBF) and metabolism. Using a porcine model of SCI, we demonstrate that high-thoracic SCI acutely impairs cardiac contractility and causes substantial reductions in intraparenchymal SCO_2_ and SCBF within the first hours post-injury. Utilizing the same model, we next show that treating the reduced contractile function with the β-agonist dobutamine is more efficacious at increasing SCO_2_ and SCBF than the current clinical standard of vasopressor therapy, whilst also mitigating increased anaerobic metabolism and hemorrhage in the injured cord. Our data provide compelling evidence that cardio-centric hemodynamic management represents a novel and advantageous alternative to the current clinical standard of vasopressor therapy for acute traumatic SCI.

## Introduction

The acute phase following a traumatic spinal cord injury (SCI) represents an important therapeutic window of opportunity to intervene with neuroprotective approaches that might limit secondary damage to the injured cord^1^, thereby providing the patient with the best chance of attaining some neurological recovery. Hemodynamic management is one of the only neuroprotective strategies available to clinicians, and current guidelines suggest that mean arterial pressure (MAP) be maintained between 85-90 mmHg with intravenous fluids and vasopressors such as norepinephrine (NE), with the aim of offsetting systemic hypotension and maintaining adequate spinal cord perfusion^2^. Though this “one-size-fits-all” strategy can improve spinal cord blood flow (SCBF), vasopressor management with NE has been shown to produce potentially harmful SCBF profiles in some acute SCI patients^3^ and has been shown by multiple investigators to exacerbate intraparenchymal hemorrhage^4–6^. In the setting of acute SCI, clinical studies have shown strong associations between increased cord hemorrhaging and worsened neurological outcomes (i.e. more neurologically complete injuries)^7^. Such hemorrhaging is therefore a critically concerning side-effect in the application of vasopressor therapy as a first-line hemodynamic management strategy for patients with acute SCI. To date, the clinical literature and therapeutic approaches have not considered that cardiac contractile dysfunction may occur acutely post-SCI and contribute to reduced spinal cord oxygenation (SCO_2_) and SCBF. As such, a hemodynamic management strategy that focuses on the heart has not been forthcoming. Only a single published study has considered the use of inotropic agents such as dobutamine (DOB) for hemodynamic management of acute SCI^8^, however the efficacy of cardiac inotropes in improving cardiovascular and spinal cord hemodynamics has not been directly compared with that of vasopressor-based management strategies that focus solely on MAP.

Accordingly, the aims of the current research were twofold. In experiment 1, we sought to define the acute impact of high-level SCI on left ventricular (LV) contractility (i.e. end-systolic elastance; E_es_) using our porcine model of contusion SCI at the second thoracic spinal cord level (T2)^9^. In experiment 2, we conducted a randomized intervention trial in the same porcine model to compare the efficacy of using the cardiac-specific β-agonist DOB versus NE (i.e., current clinical standard) in augmenting SCO_2_ and SCBF acutely following T2 SCI. To address these aims, a total of 22 female Yucatan minipigs were instrumented with a LV pressure-volume admittance catheter and pulmonary artery catheter (Fig. 1a), as well as intraparenchymal probes for SCO_2_, SCBF and microdialysis placed 1.2 cm and 3.2 cm caudal to the site of injury. Animals received a T2 contusion injury (50 g weight drop, 20 cm height) with 2 hours compression (150 g total), and hemodynamic management with DOB or NE began 30 mins post-SCI up until 4 hours post-SCI (experimental endpoint). Here, we demonstrate first that LV load-independent contractile function, including E_es_, is impaired within the first hour following a T2 SCI, and thereafter find that treating reduced contractile function with DOB is more efficacious than NE with respect to optimizing hemodynamics, improving the spinal cord microenvironment and reducing intraparenchymal hemorrhage.

**Figure 1.**
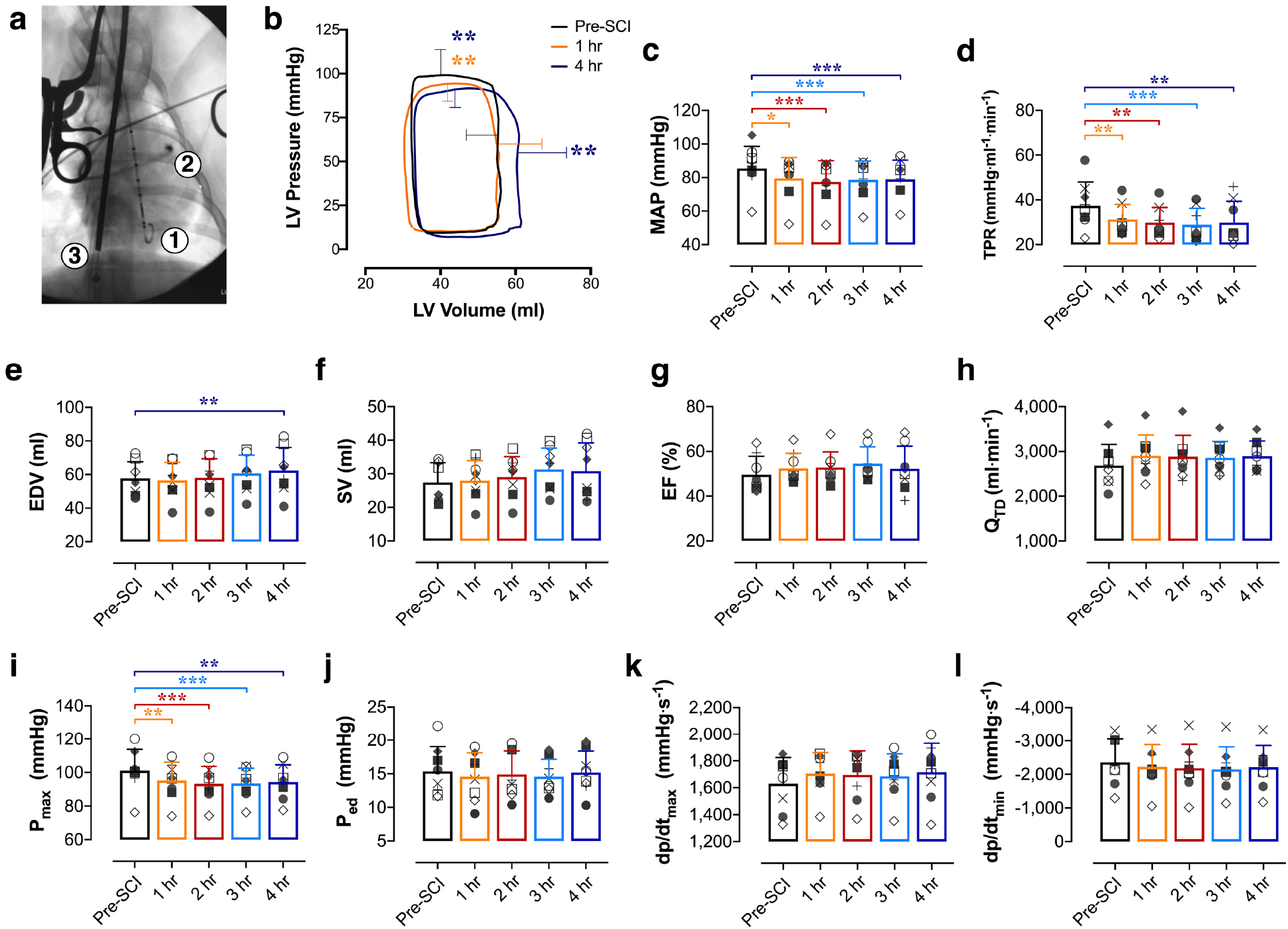
Altered load-dependent LV function and hemodynamics acutely following T2 SCI (*n=*8). (**a**) Cardiac instrumentation with a LV pressure-volume admittance catheter [1], Swan-Ganz catheter [2] advanced to the pulmonary artery and a balloon-tip inferior vena cava (IVC) occlusion catheter [3] for transient reductions to preload and assessments of LV end-systolic elastance (E_es_). (**b**) Basal pressure-volume loops represent mean interpolated data (SD bars shown for peak systolic pressure, P_max_, and end-diastolic volume, EDV) across the cardiac cycle at baseline (pre-SCI, black), 1 hour (orange) and 4 hours (blue) post-SCI. P_max_ was reduced within the first hour post-SCI and remained lowered up to 4 hours post-SCI. EDV was increased compared to pre-SCI at 4 hours, though LV stroke volume (SV) was not significantly altered by the experiment end-point. (**c**) Mean arterial pressure (MAP) and (**d**) total peripheral resistance (TPR) were reduced within the first hour post-SCI, and those reductions were sustained up to 4 hours post-SCI. (**e**) While EDV was augmented at 4 hours post-SCI, (**f**) SV, (**g**) ejection fraction (EF) and (**h**) cardiac output (Q_td_) were unchanged post-SCI. (**i**) P_max_ was impaired within 1 hour post-SCI, but (**j**) end-diastolic pressure (P_ed_), (**k**) the maximal rates of systolic pressure generation (dp/dt_max_) and (**l**) diastolic pressure decay (dp/dt_min_) were unchanged within 4 hours post-SCI. *p<0.05, **p<0.01, ***p<0.001 versus pre-SCI. Data represent means ± standard deviation (SD), and were assessed using a one-way repeated measures ANOVA with Fisher’s LSD for post-hoc comparisons versus pre-SCI data.

## Results

### Cardiac contractility is impaired in acute T2 SCI

In experiment 1 (*n*=8), LV maximal systolic pressure (P_max_), MAP and total peripheral resistance (TPR) were reduced within 1 hour post-SCI, and remained depressed up to 4 hours post-SCI (Fig. 1b, Fig. 1c and Fig. 1d). At 4 hours post-injury there was a slight but significant increase to LV filling volume (i.e. EDV; Fig. 1e); however, there were no significant alterations to LV stroke volume (SV, Fig. 1f), ejection fraction (EF, Fig. 1g) or cardiac output measured with thermodilution (Q_TD_, Fig. 1h) within the 4 hours following T2 SCI.

The major finding from experiment 1 was that LV contractility was immediately impaired within the first hour post-SCI. Specifically, we observed that LV load-independent systolic function assessed as E_es_ (Fig. 2a and Fig. 2b), preload-recruitable stroke work (Fig. 2d) and maximal rates of pressure generation for a given filling volume (Fig. 2e) were all reduced by 1 hour post-SCI and remained depressed until 4 hours post-SCI. We additionally examined LV contractile reserve utilizing a high-dose DOB challenge (i.e. 10 μg/kg/min DOB) before and 4 hours following T2 SCI, and found that contractile reserve was compromised post-SCI compared to baseline (Fig. 2c). In contrast to the clear impairments to LV systolic function, LV load-independent diastolic function as assessed with the end-diastolic pressure-volume relationship (EDPVR) was not altered acutely post-SCI (Fig. 2f). Measures of load-dependent diastolic function, including LV end-diastolic pressure (P_ed_) and the rates of diastolic pressure decay (-dp/dt_min_ and **τ**), were also unaltered from baseline in the 4 hours following injury (Supplemental Table S1a).

**Figure 2.**
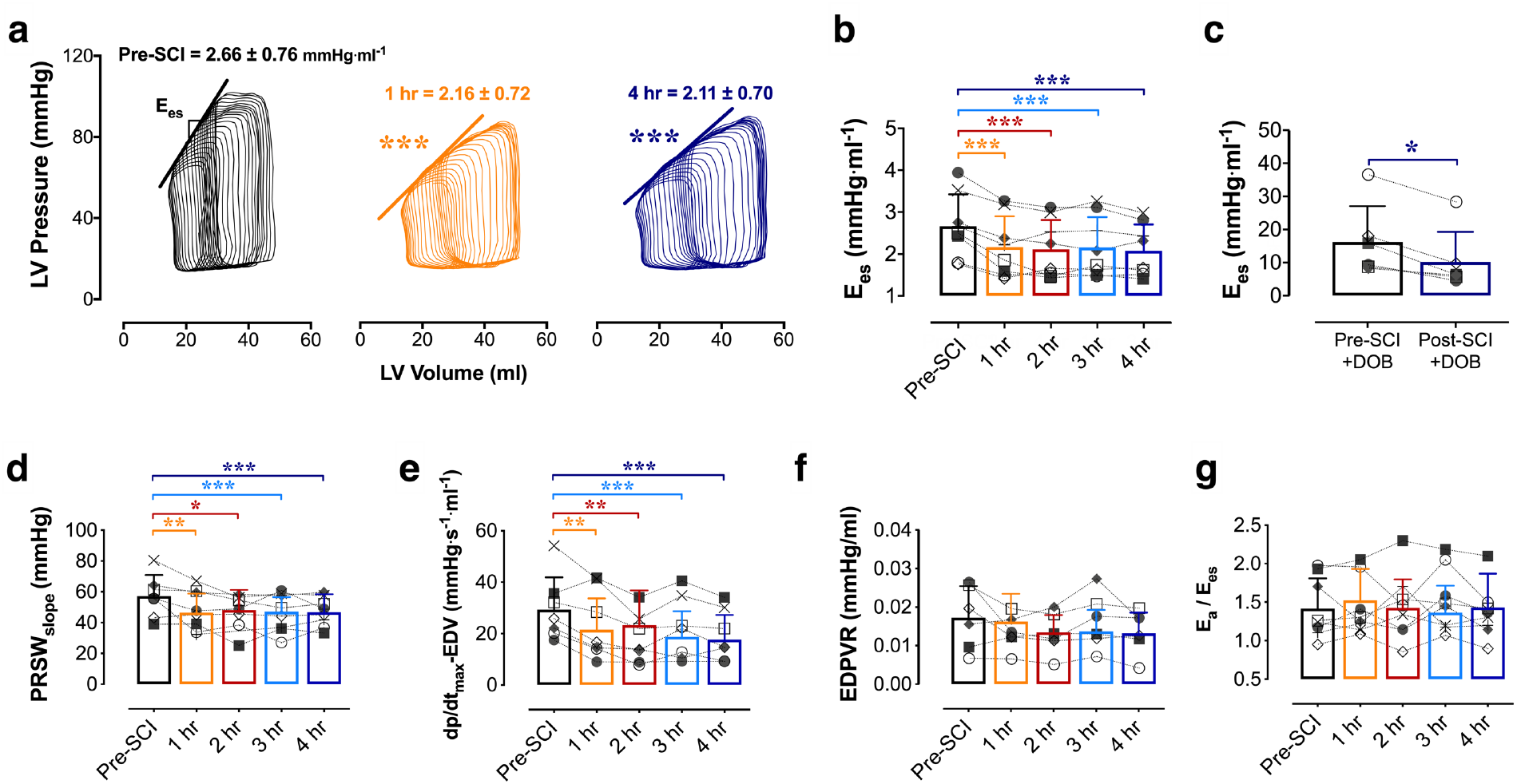
Impaired LV systolic load-independent function post-SCI. (**a**) Representative PV loops during IVC occlusions illustrating impaired LV contractility (end-systolic elastance; E_es_) within 1 hour (middle, orange) and up to 4 hours (right, blue) post-SCI. (**b**) Group data showing the reduction to E_es_ acutely post-SCI. (**c**) Animals additionally had reduced E_es_ responses to dobutamine challenges (‘DOB’, 10 μg/kg/min) post-SCI. (**d**) Impaired systolic load-independent function is further supported by the early (1 hour) and sustained reductions to preload-recruitable stroke work (PRSW) and (**e**) the rate of maximal pressure generation for a given EDV (dp/dt_max_-EDV). (**f**) The end-diastolic pressure volume relationship (EDPVR) was not altered acutely post-SCI. (**g**) There were no changes to the relationship of arterial elastance (E_a_) to E_es_, due to simultaneous reductions in LV afterload and contractility following SCI. Statistics are identical to those outlined in Figure 1.

### High-dose DOB optimizes LV function and hemodynamics

In experiment 2, we utilized a randomized and counterbalanced design whereby 14 additional animals (*n=*7 per group) received individualized hemodynamic management with either DOB or NE starting 30 minutes post-SCI (Fig. 3a). Drug levels were continually titrated to achieve a target E_es_ of ~2.5-2.9mmHg/ml for animals receiving DOB, based on the baseline pre-SCI mean E_es_ from animals in experiment 1, and a target MAP of ~85-90mmHg for animal receiving NE, in line with the current clinical guidelines^2^. As a result of this individualized approach, 4 of the animals receiving DOB were administered higher doses (i.e., ≥2.5 μg/kg/min, DOB+) while 3 received negligible doses (i.e., ≤0.5 μg/kg/min, DOB-; Fig 3a). As such, we subsequently stratified the DOB animals by dose (i.e. DOB+ and DOB-groups). All NE animals received sufficient doses to maintain MAP at 85-90mmHg (mean 0.16μg/kg/min, range 0.06-0.46μg/kg/min), which were similar to those reported in our group’s previous studies using a low-thoracic SCI porcine model^10^.

**Figure 3.**
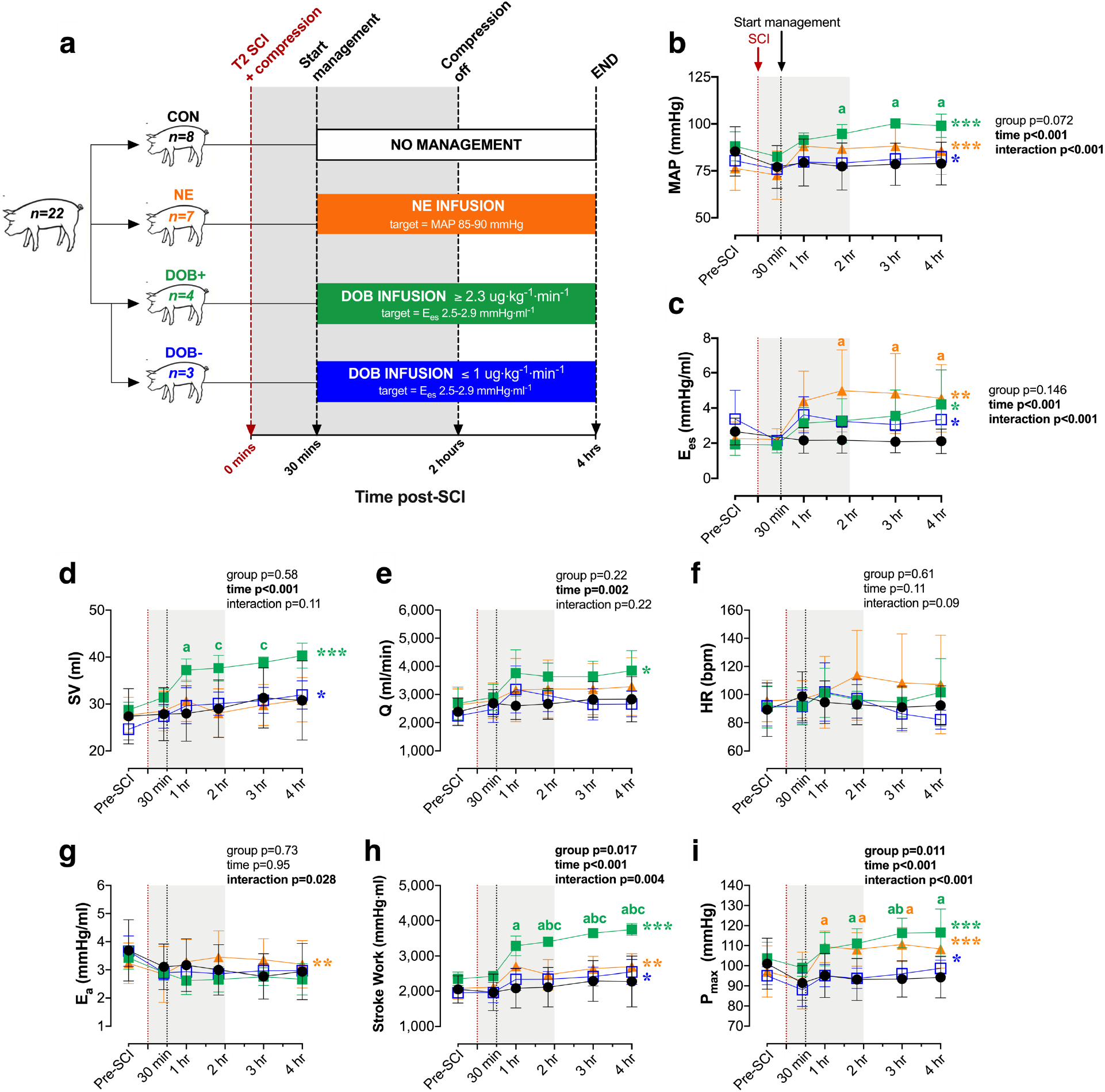
Impacts of hemodynamic management with dobutamine (DOB) and norepinephrine (NE) on hemodynamics and LV function following acute T2 SCI. Dependent variables from 30 mins (i.e. start of hemodynamic management) to 4 hours post-SCI were analyzed using a two-way repeated-measures ANOVA (factors: time, group), and post-hoc comparisons were made with Tukey’s test (between-group) and Fisher’s LSD (within-group compared to 30 mins). (**a**) Following experiment 1 (control animals, CON, black), an additional 14 animals received hemodynamic management starting 30 mins post-SCI with either DOB titrated to a target E_es_ of ~2.5-2.9 mmHg/ml, or NE (orange) titrated to a mean arterial pressure (MAP) of 85-90 mmHg. Three animals receiving DOB had a relatively high baseline E_es_, and as a result received minimal doses of DOB (i.e., ≤0.5μg/kg/min, DOB-, blue) while 4 animals received substantial DOB doses (i.e., ≥2.5μg/kg/min, DOB+, green). (**b**) Both DOB+ and NE augmented MAP, however MAP was significantly augmented in DOB+ compared to CON animals. (**c**) LV contractility, E_es_, was augmented by both DOB and NE. (**d**) Only DOB+ generated increases in LV stroke volume (SV) and (**e**) cardiac output (Q), which were not observed with NE. (**f**) There were no significant changes to heart rate (HR) with hemodynamic management, nor were there differences between the groups. (**g**) LV afterload increased with NE management but not with DOB. (**h**) Stroke work, an index of systolic function, is only increased with DOB+, (**i**) despite increases to P_max_ in both NE and DOB. *p<0.05, **p<0.01, ***p<0.001 (within-group effect for time). ^**a**^p<0.05 vs CON; ^**b**^p<0.05 vs DOB -; ^**c**^p<0.05 vs NE.

Hemodynamic management with DOB+ and NE both augmented MAP and LV contractility (E_es_) up to 4 hours post-SCI (Fig. 3b and Fig. 3c); however, the two drugs achieved increases to MAP via markedly different alterations to cardiac and vascular hemodynamics. Specifically, DOB+ increased MAP via improvements in LV systolic function (Fig. 3d and Fig. 3h) and cardiac output (Fig 3e), whereas NE restricted stroke volume and cardiac output (Fig. 3d and Fig. 3e) by significantly augmenting LV afterload (E_a_, Fig. 3g). DOB+ additionally enhanced LV early diastolic relaxation (**τ**, Supplemental Table S2b), which was not observed with NE despite both groups having similar heart rates throughout the experiments (Fig. 3f). DOB-animals demonstrated small but significant improvements in LV contractility (E_es_), stoke work, stroke volume and MAP, however due to the very small doses of DOB received by DOB-animals those hemodynamic effects were minimal in comparison to DOB+.

### High-dose DOB improves spinal cord oxygenation and mitigates cord hemorrhaging

Within the spinal cord parenchyma, DOB+ animals exhibited large improvements to SCO_2_ (measured at the 1.2 cm probe) following decompression (Fig. 4b and Fig. 4c), and the relative increase to SCO_2_ was greatest in DOB+ compared to both CON and NE animals at 3 hours (p=0.05 vs. NE; p=0.02 vs. CON) and 4 hours post-SCI (p=0.02 vs. NE and CON). During the compression period (i.e., initial 2 hours post-SCI), only DOB+ appeared to improve SCBF (p=0.028 vs. CON at 2 hours post-SCI; Fig. 4d), while SCBF remained depressed in all other animals. DOB+ additionally mitigated increases in the lactate-to-pyruvate ratio during the compression period (Fig. 4f) that were otherwise observed in the NE and CON animals.

**Figure 4.**
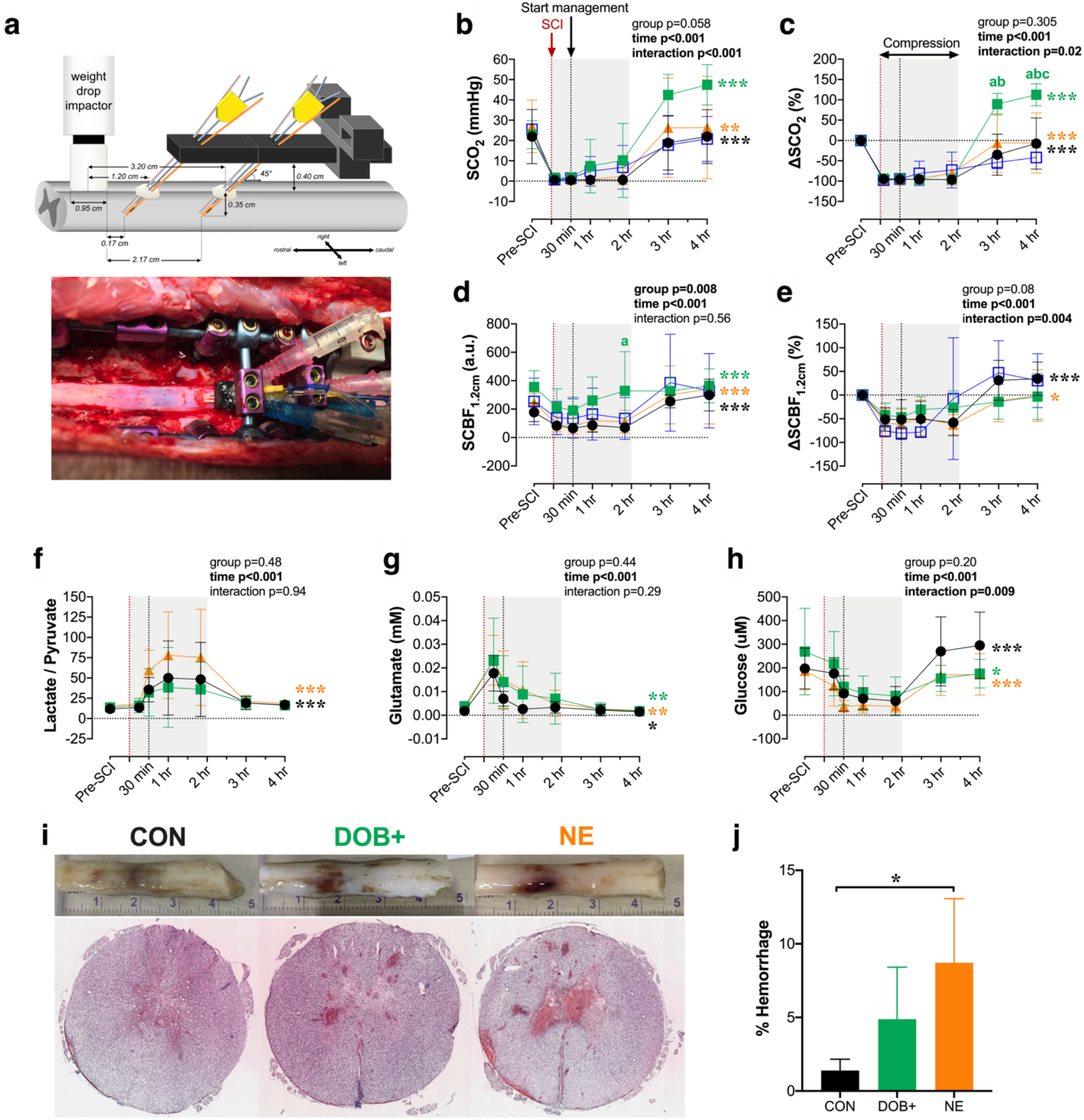
Impact of hemodynamic management on spinal cord oxygenation (SCO_2_), blood flow (SCBF), metabolism and hemorrhage acutely post-SCI. (**a**) Setup for intraparenchymal monitoring. A fixation device is secured to the spinal column and probes are inserted through the dura 1.2 cm and 3.2 cm away from the center of the impactor. All data are shown for 1.2 cm probes. Data from 3.2 cm probes are provided in Supplemental Table S2d. (**b**) Improvements to SCO_2_ occur after decompression (i.e. 2 hours post-SCI), but are most pronounced in DOB+. (**c**) When expressed as a percent change form baseline, ΔSCO_2_ is significantly augmented in DOB+ compared to all groups by 4 hours post-SCI. (**d**) SCBF is augmented in CON, NE and DOB+ following decompression, and DOB+ notably had augmented absolute SCBF at 2 hours post-SCI. (**e**) However, when expressed as percent change from baseline, SCBF was only altered in CON and NE over the treatment period. (**f**) The lactate/pyruvate ratio does not increase after management onset (i.e. 30 mins post-SCI) with DOB+, but is increased in NE and CON. (**g**) In all groups, glutamate becomes progressively reduced and (**h**) glucose is increased following decompression (i.e. 2 hours post-SCI). Sufficient microdialysis data were only acquired in *n=*2 for DOB-, thus DOB-data were excluded from analyses. (**i**) Representative cords and histological stains show pronounced hemorrhaging with NE. (**j**) Animals receiving NE have augmented hemorrhaging at the injury epicentre, which is mitigated by DOB+. See Figure 3 for statistical analyses, additional abbreviations and colour legends. *p<0.05, **p<0.01, ***p<0.001. ^**a**^p<0.05 vs CON; ^**b**^p<0.05 vs DOB-; ^**c**^p<0.05 vs NE.

Histological analyses at the injury epicenter demonstrated that NE exacerbated spinal cord hemorrhaging compared to CON (Fig. 4i and Fig. 4j), whereas animals receiving DOB+ were spared the increased hemorrhaging. Immunohistochemical analyses of the injury epicentre did not reveal any between-group differences in the densities of glial fibrillary acidic protein (GFAP+) or inflammatory activation (IBA1+, Fig. 5).

**Figure 5.**
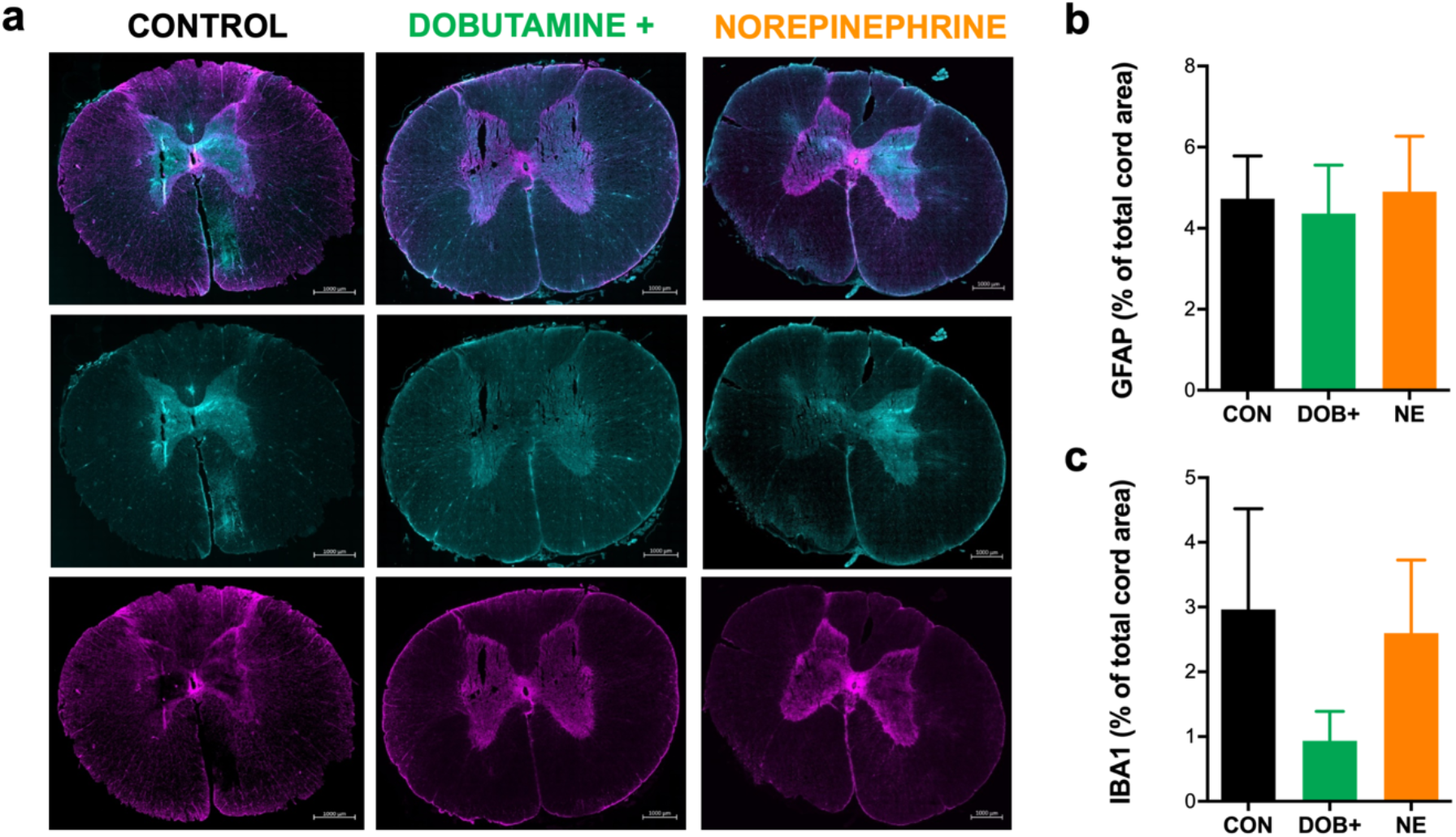
Immunohistochemical analyses of glial (GFAP+) and inflammatory activation (IBA1+) in the acutely injured spinal cord epicentre. (**a**) Representative images are shown for control (left) animals, and animals receiving hemodynamic management with high-dose dobutamine (middle) and norepinephrine (right). Merged stains (top) are shown for ionized calcium binding adaptor molecule 1 (IBA1+, bottom) and glial fibrillary acidic protein (GFAP+, middle). Group data are shown for immunohistochemical analyses of IBA1 **(b)** and GFAP **(c).** Bars represent means ± standard deviations (SD). No significant differences were detected between animals in control (CON), high-dose dobutamine (DOB+) and norepinephrine (NE) groups. *n=*4 per group for immunohistochemical analyses.

## Discussion

Our findings provide compelling evidence that LV load-independent contractility is immediately impaired in acute high-level SCI, and that correcting LV contractility with DOB+ is beneficial to the spinal cord parenchyma by optimizing cord oxygenation and blood flow via cardio-genic improvements in MAP absent of systemic vasoconstriction. Furthermore, our data highlight that the current clinical standard of hemodynamic management with NE does not support improved LV function, does not modify SCBF and in fact appears to worsen hemorrhaging. This research therefore supports the efficacy of implementing a cardiac-focused hemodynamic management strategy in the acute phase following high-thoracic SCI.

In experiment 1, the immediate reductions to key load-independent measures of LV systolic function incontrovertibly indicates that intrinsic contractile dysfunction in high-level SCI results from the immediate loss of descending sympathetic input following high-level injury. Previously, only a small collection of echocardiographic studies in humans had provided some evidence for chronically-altered LV systolic function in humans^11^, and the interpretation of those findings were limited due to the load-dependent nature of echo-derived data. To assess load-independent LV function our group has utilized LV pressure-volume catheterization in a chronic rodent model of SCI and reported reductions to LV E_es_ after a T2 injury^12,13^. Our present data extend these observations from the chronic setting by demonstrating that LV contractility is impaired immediately following high-level SCI. Importantly, we highlight that EF was unchanged despite clear reductions to LV contractility, reinforcing that EF does not adequately detect systolic or contractile dysfunction in SCI^11^. We have additionally identified a reduction to LV contractile reserve acutely after the injury, which may be attributable to a rapid loss of ‘baseline’ contractility and tonic neuro-hormonal activation of the myocardium that occurs following high-level SCI. Though our group has previously reported that systolic reserve is relatively intact in chronically-injured rats with T2 SCI^13,14^, those observations may reflect chronic hyper-responsiveness of cardiac β-adrenergic receptors^15,16^, which would not have occurred in the acute setting of the current study.

The impacts of acute high-level SCI on LV diastolic function are less clear, given there were no significant alterations to diastolic indices within the 4 hours following T2 SCI. The lack of changes to the EDPVR (load-independent measure of LV compliance or stiffness) and load-dependent diastolic pressure decay is presumably a function of time, given that long-term structural remodelling and tissue stiffening generally precede diastolic dysfunction^17^. While there was a small increase to EDV by 4 hours post-SCI, this may be explained by re-lengthening of the cardiomyocytes following the loss of tonic sympathetically-mediated β-adrenergic activation^18^. The impacts of SCI on the diastolic phase are not well-characterized amongst the literature, though some pre-clinical work from our group has identified blunted relaxation rates^12^ alongside significant myocardial fibrosis in rodents with chronic high-level SCI^19^, underscoring the time-dependency of altered diastolic function. Nonetheless our current data provide new evidence that diastolic function is not critically impacted within the first hours following high-level SCI.

In experiment 2, we demonstrate that correcting LV contractility with higher doses of DOB (i.e. DOB+) optimized both cardiac and spinal cord outcomes acutely post-SCI more effectively than the current clinical standard of hemodynamic management with vasopressors (i.e. NE). Though both approaches effectively augmented MAP, this was achieved with DOB+ by increasing cardiac output, whereas NE predominantly increased vascular resistance and LV afterload (E_a_) precluding any improvements to LV systolic output. This substantial arterial afterload is concerning as it may additionally stress or damage the myocardium if management is prolonged, and further exacerbate the long-term negative consequences of SCI on the heart^11^.

Within the spinal cord microenvironment, DOB+ appeared to alleviate spinal cord ischemia more effectively than NE by optimizing cord oxygenation, blood flow and metabolic indices. In contrast, NE did not modify SCBF and worsened hemorrhaging, mirroring observations from Soubeyrand et al. ^4^ in a feline model. Several studies have linked NE with central gray matter hemorrhaging in experimental models^5,6^, which is thought to result from unfavourable blood flow redistribution in the cord microenvironment^3^. Specifically, NE may reduce flow in the intact cord circulation via α_2_-mediated vasoconstriction^20,21^ and subsequently worsen blood loss and hemorrhage through the damaged microvasculature. In contrast to NE, DOB does not directly alter vascular tone, and we contend it may facilitate improved shear-mediated vasodilation ^22^ of the cord vasculature via increases in cardiac output, rather than augmenting the driving pressure to the site of hemorrhage. Collectively, our data provide compelling evidence that DOB+ has a beneficial effect on the spinal cord parenchyma via cardio-genic improvements in MAP absent of systemic vasoconstriction.

This research supports the efficacy of implementing a cardiac-focused hemodynamic management strategy in the acute phase following high-thoracic SCI. We have demonstrated that correcting the reduction in LV contractility with higher doses of DOB optimizes LV function, hemodynamics and local cord oxygenation more effectively than the current clinical standard of NE despite similar elevations to MAP. We therefore contend that the method by which MAP is elevated has a profound effect on the injury site microenvironment. By non-selectively binding both β-and α-adrenergic receptors, NE produces systemic vasoconstriction to augment MAP but at the cost of large increases to cardiac afterload which oppose any potential improvements to cardiac output. We therefore conclude that a cardiac-specific strategy provides a more advantageous approach for optimizing hemodynamic management in acute SCI than standard vasopressor treatment. These findings merit clinical investigation, given that hemodynamic management is one of the few neuroprotective treatment options for acute SCI patients.

## Materials and Methods

### Ethics, animals and handling

All protocols and procedures were compliant with Canadian Council on Animal Care policies, and ethical approval was obtained from the University of British Columbia Animal Care Committee (A16-0311) and United States Department of Defence (IACUC #A16-0311).

22 female Yucatan minipigs aged 2-3 months (20-25 kg; S&S Farms, Ramona, CA, USA) were acquired and housed in the Centre for Comparative Medicine (University of British Columbia, South Campus) animal facility for 1-2 weeks prior to surgery. Animals were housed in pairs or small groups (4-6 animals) in indoor pens with sawdust bedding and with access to an adjoining outdoor pen. Animals received daily visits from a researcher to become habituated to humans, were provided enrichment (toys e.g. chains, balls), water ad libitum and feed equal to 1.5% of body mass twice per day.

### Experimental overview

#### Experiments 1 and 2

All animals were instrumented similarly for cardiovascular and spinal cord measurements, with bilateral pressure catheters in the femoral arteries, a pressure-volume (PV) admittance catheter placed in the left ventricle (LV), a Swan-Ganz thermodilution catheter placed in the pulmonary artery and a venous balloon occlusion catheter advanced to the inferior vena cava (IVC) via the right femoral vein. The PV and Swan-Ganz catheters were advanced under fluoroscopic guidance. Placement was confirmed via the emergence of a typical pressure-volume loop and typical pulmonary artery pressure waveforms. A laminectomy was then performed to expose the spinal cord from the C8-T4 level, and custom-designed sensors for spinal cord blood flow, oxygenation, pressure, and microdialysis were placed in the spinal cord parenchyma at 1.2 cm and 3.2 cm caudal to the impactor centre^23^. Once drug levels and sensors had stabilized (approximately 2-3 hours after laminectomy), baseline data for cardiac function, hemodynamics and spinal cord indices were obtained over a 30-minute period. Following baseline data collection, animals received a T2 weight-drop (50 g) contusion injury with 2 hours of spinal cord compression (additional 100 g, 150 g total). Cardiac, hemodynamic and spinal cord indices were continuously recorded up to 4 hours post-SCI, at which point animals were euthanized and spinal cord tissue was immediately harvested.

#### Experiment 1: Effect of SCI on contractile function and reserve (n=8)

Once a stable plane of anaesthesia was reached prior to SCI, and hourly post-SCI, contractile function was assessed with transient IVC occlusions for characterization of load-independent systolic function, including end-systolic elastance (E_es_), preload-recruitable stroke work (PRSW) and the maximal rate of pressure generation for a given end-diastolic volume (dp/dt_max_-EDV). Contractile reserve was assessed using a constant-rate infusion of DOB (10 μg/kg/min) via an infusate port in the Swan-Ganz catheter for 10 minutes. This dosage has been previously utilized to challenge LV load-dependent^24^ and contractile function in pigs^25^. Within the final 2 minutes of DOB infusion, LV load-independent contractility was assessed with transient IVC occlusions for characterization of the E_es_. A minimum of 30 minutes recovery (≥5 half-lives of DOB^26^) was provided before ‘baseline’ measurements began pre-SCI, and before euthanasia and tissue collection at the end of the experiments.

#### Experiment 2: Comparing post-SCI hemodynamic management strategies (n=7 per group)

For animals receiving hemodynamic management with NE or DOB, treatment type was randomized and counterbalanced between groups. Drugs were administered via an infusion port on the Swan Ganz catheter, beginning at 30 minutes post-SCI until 4 hours post-SCI. Infusions of NE were titrated to attain a target MAP of 85-90 mmHg^8^, and DOB was titrated to attain an E_es_ slope of ~2.5-2.9 mmHg/ml, based on the average pre-SCI E_es_ slope observed in experiment 1. Drugs were titrated continually through the first 30 minutes of infusion to attain the given target, and hourly thereafter until 4 hours post-SCI.

### Specific Methodology

#### Surgical preparation and anaesthesia

Animals were fasted for 12 hours prior to surgery, pre-anaesthetized with intramuscular injections of telazol (4-6 mg/kg), xylazine (1 mg/kg) and atropine (0.02-0.04 mg/kg), and thereafter induced with propofol (2 mg/kg). Animals received endotracheal intubation for mechanical ventilation (10-12 breaths/min; tidal volume 12-15 ml/kg; Veterinary Anesthesia Ventilator model 2002, Hallowell EMC, Pittsfield, MA). A urinary catheter (10 F, Jorgensen Laboratories Inc., Loveland, CO) was placed for intra-operative bladder drainage, and intravenous catheters placed for administration of anesthetic agents and fluids. A rectal temperature probe was additional placed, and core body temperature maintained at 38.5-39.5 °C with a heating pad (T/Pump, Gaymar Industries, Inc., Orchard Park, NY). Throughout surgery, animals received intravenous continuous rate infusions of propofol (9-13 mg/kg/hr), fentanyl (10-15 mg/kg/hr) and ketamine (5-8 mg/kg/hr), as well as intravenous fluid to maintain hydration (7-10 ml/kg/hr, 2.5% dextrose + 0.9% NaCl). The surgical plane of anesthesia was determined by the absence of jaw tone assessed by the veterinarian technicians. Standard monitoring was performed for heart rate (electrocardiogram) respiratory rate, end tidal carbon dioxide, MAP, and oxygen saturation (pulse oximeter 8600V, Nonin Medical Inc., Markham, ON).

#### Cardiac and arterial catheterization

After induction, animals were transferred to an operating table and oriented in a supine position. Five-centimeter incisions were made on the medial side of both hindlimbs, and tissue bluntly dissected to reveal the femoral arteries. Two-inch intravenous catheters (20 g) were advanced into the arteries and connected to fluid-filled lines. Amplifiers and pressure transducers connected the arterial lines to an A/D board (PowerLab, ADInstruments, Colorado Springs, MO) for real-time monitoring for blood pressure (i.e., systolic blood pressure [SBP], diastolic blood pressure [DBP] and MAP) with commercially-available software (LabChart PRO v8.1.9, ADInstruments). Two catheters were utilized in case one of the lines failed while repositioning the animal to the prone position.

For placement of cardiac catheters, a 5 cm incision was made in the tissue overlaying the right jugular vein, and blunt dissection revealed the carotid artery and external jugular vein. Prior to insertion, channels for the admittance PV catheter (5F; Sciscence Catheter and ADVantage PV System [ADV500], Transonic Systems Inc.) and Swan-Ganz thermodilution catheter (7.5 F; Edwards Lifesciences Canada Inc., Mississauga, ON) were connected to the A/D board for real-time visualization of catheter pressures. The pigtailed PV catheter was inserted with an introducer (12 F; Fast-Cath Hemostasis Introducer, Abbott) into the carotid artery and advanced until an arterial waveform was visualized, and the Swan-Ganz catheter inserted into the external jugular vein and advanced until a right ventricular pressure waveform became apparent. Both catheters were then further advanced under combined pressure and fluoroscopic guidance (Arcadis Avantic, Siemens Healthcare Limited, Oakville, ON) to ultimately place the PV catheter into the LV and the Swan-Ganz catheter into the pulmonary artery. Sutures were placed around the vessels and catheters to secure placement and the tissue was closed.

#### Laminectomy

Following placement of arterial and cardiac catheters, animals were reoriented to the prone position, and the spinous processes, laminae and transverse processes of the C8-T7 spine were exposed with electrocautery. Using anatomical landmarks, two 3.5×24 mm multi-axial screws (Select™ Multi Axial Screw, Medtronic, Minneapolis, MN) were placed into the T1 and T4 pedicles. A 3.2 mm diameter titanium rod (Medtronic, Minneapolis, MN) was affixed to the screws to rigidly fix the T1-T2-T3 segments and additionally secure the weight drop system. A T2-T3 laminectomy was performed to provide a circular window ≥1.2 cm in diameter exposing the dura mater and spinal cord, then the C8-T4 laminae were further resected to expose the spinal cord and allow for insertion of sensors and catheters surrounding the injury site.

#### Implantation of spinal cord blood flow, oxygenation, pressure and microdialysis probes

Probes for intraparenchymal spinal cord monitoring and microdialysis were placed as previously described^10^. Briefly, a custom-made sensor platform was secured over the titanium rods and adjusting the pedicle screws. Six custom introducers were inserted through the platform at 45° angles, entering the dura at 1.2 cm and 3.2 cm caudal to the injury epicenter. Sensors for blood flow and oxygenation (combined), pressure and microdialysis were guided through the introducers into the ventral aspect of the white mater, with final placement of the catheter tip centers ~2 mm (proximal probes) and 22 mm (distal probes) from the edge of the impactor. Placement in the spinal cord was confirmed with a commercially-available ultrasound system (LOGIQ e Vet, GE Healthcare, Fairfield, CT) using a linear array 4-10 MHz transducer (8L-RS). Cyanoacrylate glue was applied to the dural surface surrounding catheter implantation to prevent cerebrospinal fluid leakage. A minimum of 2 hours was provided for intraparenchymal probe stabilization prior to the collection of baseline data.

#### Spinal cord injury

A weight-drop impactor device with an articulating arm (660, Starrett, Athol, MA) and guide rail was mounted on a metal base and secured to the T1 and T4 vertebra with the pedicle screws described above. The tip of the impactor (0.953 cm diameter), equipped with a load cell (LLB215, Futek Advanced Sensor Technology, Irvine, CA) to acquire force of impact data, was oriented vertically above the exposed dura and cord at the T2 level. The guide rail was equipped with a linear position sensor (Balluff Micropulse®, Balluff Canada Inc., Mississauga, ON) to obtain data on impactor position during the weight drop. The device was remotely operated using LabVIEW software (National Instruments, Austin, TX), which additionally acquired real-time impactor force and position data. Five minutes prior to injury, animals received a continuous-rate infusion of fentanyl (7 μg/kg over 1 minute). The SCI was carried out by dropping a 50 g cylinder plastic weight through the guide rail from a height of approximately 16 cm, with another 100 g weight added immediately following the initial weight-drop for a total 150 g compression. At 2 hours post-SCI, the compression weight and spinal cord injury device were removed (decompression), after which pedicle screws were removed and bone wax was used to close screw holes in the vertebral body.

#### Measurement and analysis of load-dependent and load-independent LV function

During experiments, LV pressure and volume data were continuously obtained from the PV catheter, as outlined above, and all analyses of LV-PV data were performed off-line using LabChart PRO software (v8.1.9, ADInstruments, Colorado Springs, CO) with the PV Loop Analysis module. Load-dependent measures of LV pressure indices (maximal pressure [P_max_], minimum pressure [P_min_], end-systolic pressure [P_es_], end-diastolic pressure [P_ed_], maximum rate of pressure generation [d_p_/d_tmax_], maximal rate of pressure decay [d_p_/d_tmin_], time constant of diastolic pressure decay [τ]), volumetric indices (end-diastolic volume [EDV], end-systolic volume [ESV], stroke volume [SV], ejection fraction [EF]), and stroke work and arterial elastance (E_a_) were assessed from basal loops over a 1-minute period immediately preceding the defined measurement point (i.e. prior to IVC occlusions and thermodilution).

For the assessment of load-independent LV function, LV preload was manipulated using transient IVC occlusions at baseline (pre-SCI), 30 minutes post-SCI (just prior to hemodynamic management in experiment 2) and then hourly up to 4 hours post-SCI. In experiment 1, IVC occlusions were also performed during the DOB challenge. Analysis of approximately 10-15 PV loops during IVC occlusions allowed for assessments of load-independent systolic and diastolic function, including E_es_ and the end-diastolic pressure-volume relationship (EDPVR), respectively.

#### Measurement of pulmonary pressures and cardiac output via Swan-Ganz catheterization

During experiments, pulmonary artery pressures were monitored in real-time and continually recorded from the Swan-Ganz catheter. Cardiac output (Q) was measured with the thermodilution technique at baseline (pre-SCI), 30 minutes (prior to hemodynamic management) and then hourly post-SCI. Briefly, bolus infusions (~10 ml) of iced saline (0-6 °C) were administered through a temperature-recording flow-through housing (REF: 93505, Edwards Lifesciences Canada Inc., Mississauga, ON) and into the proximal port of the Swan-Ganz catheter. Bolus infusions for thermodilution were always performed ≥1 minute following IVC occlusions. Calculations of Q were performed off-line using LabChart PRO software (v8.1.9, ADInstruments, Colorado Springs, CO) with the Cardiac Output Analysis module (v1.3).

#### Measurement of spinal cord oxygenation, blood flow and pressure, and metabolism

Measurements of spinal cord oxygenation and blood flow were obtained in real-time using a multi-parameter probe (NX-BF/OF/E, Oxford Optronix, Oxford, UK) attached to a combined OxyLab/OxyFlo channel monitor, as previously described^10^. Spinal cord partial pressure of oxygen (SCO_2_) was monitored with fiber optic oxygen sensors that utilize the fluorescence quenching technique^27^. Relative changes in SCBF were monitored with laser-Doppler flowmetry. Spinal cord pressure was assessed with custom-manufactured fiber optic pressure transducers (FOP-LS-NS-1006A, FISO Technologies Inc., Harvard Apparatus, Quebec, Canada) that employ Fabry-Pérot interferometry^10,28^. Pressure transducers were connected to a signal conditioner module (EVO-SD-5/FPI-LS-10, FISO Technologies Inc., Harvard Apparatus, Quebec, Canada) and data were continually recorded using the Evolution software (FISO Technologies Inc., Harvard Apparatus, Quebec, Canada).

Energy-related metabolites were measured with microdialysis probes (CMA11, CMA Microdialysis, Harvard Apparatus, Quebec, Canada) as outlined previously^10^. Briefly, a subcutaneous implantable micropump (SMP-200, IPrecio, Alzet Osmotic Pumps, Durect Corporation, Cupertino, CA) was used to continuously perfuse probes with artificial cerebrospinal fluid (Perfusion Fluid CNS, CMA Microdialysis, Harvard Apparatus, Quebec, Canada) and dialysates were acquired and frozen with dry ice every 15 minutes, starting at baseline (pre-SCI) until 4 hours post-SCI. Samples were analyzed for five metabolites (i.e., lactate, pyruvate, glucose, glutamate, and glycerol) within one week of collection (ISCUSflex Microdialysis Analyzer, M Dialysis, Stockholm, Sweden).

Measures of SCO_2_, SCBF, spinal cord pressure and microdialysis were acquired from both locations (i.e., 1.2 cm and 3.2 cm caudal to the impactor) throughout experiments 1 and 2.

#### Spinal cord tissue processing, histology and immunochemistry

Following euthanasia, 6 cm of spinal tissue surrounding the injury epicenter was removed and placed in 4% paraformaldehyde. Over the next 15 days, tissue was placed in increasing concentrations of sucrose until a concentration of 30% was reached. The tissue was then cut into 1 cm sections (ensuring the injury epicenter is within a single section), then embedded in optimal cutting temperature matrix (Shandon Cryomatrix, Thermo Scientific), frozen, and kept at −80° C. The injury section was further cut into 30 μm sections and mounted onto a series of 10 slides coated with Silane Surgipath Solution (Leica). These slides were then stored at −80° C. For histology and immunochemistry, sections were thawed at room temperature for 1 hour, at which time a hydrophobic barrier was drawn using ImmEDGE Hydrophobic Barrier Pen (Vector Laboratories). Sections were then rehydrated in 0.1 M PBS for 10 minutes then incubated with 10% normal donkey serum 0.2% Triton X-100 plus 0.1% sodium azide in PBS. Sections were incubated overnight with primary antibody rabbit anti-IBA1 (1:1000, Novus NBP2-19019). Sections were incubated for 2 hours with secondary antibody Alexa Fluor 488 donkey anti-rabbit (1:1000 abcam ab150073), and glial fibrillary acidic protein (GFAP) conjugated Cy3 produced in mouse (Sigma, C9205). Slides were cover-slipped with ProLong Gold with DAPI (Invitrogen). For hematoxylin and eosin (H&E, Leica Biosystems, San Diego, CA, USA) staining, slides were thawed at room temperature and staining was conducted using standard techniques laid out by Leica. Slides were cover-slipped with ProLong Gold (Invitrogen).

#### Analyses of immunochemistry and histology data

Immunohistochemical staining was imaged using a Zeiss Axiophot microscope (Carl Zeiss, Oberkochen, Germany) equipped with a digital camera (Olympus Q5). H&E stains were imaged using a Leica Aperio CS2 scanner (Leica Biosystems, San Diego, CA, USA). All images were processed and analyzed by standard densitometric analyses using ImageJ (U. S. National Institutes of Health, Bethesda, Maryland, USA). Briefly, quantification of the immunostaining, GFAP and IBA1, were carried out by measuring the immunopositive areas in the spinal cord section. Similarly, quantification of the H&E staining was carried out by manually outlining the area of the regions of hemorrhaging, which were identified by areas that exhibited dense red staining. All positive stains are expressed as a % of total spinal cord area. Reported values reflect means of 5 separate sections per animal.

### Statistical analysis and sample size calculation

Data are presented as means ± standard deviation (SD) in figures and supplemental tables (for non-normally distributed data, medians and interquartile ranges are provided in Supplemental Table 3). Normalcy was determined using the Shapiro-Wilk test. For experiment 1, normally distributed dependent variables were analyzed using a one-way repeated-measures analysis of variance (ANOVA). Post hoc pairwise comparisons were made with Fisher’s LSD for planned within-group comparisons, and Tukey’s HSD for between-group comparisons. Paired t-tests were used to compare data between DOB challenges pre- and post-SCI. For experiment 2, normally-distributed dependent variables were analyzed with a repeated measures ANOVA with two independent factors (group × time), and when a significant effect was detected post hoc comparisons were performed for within-group data with Fisher’s LSD, and for between-group data with Tukey’s test. For non-normally distributed data, a Friedman repeated-measures ANOVA on ranks was used to detect within-group differences over time, and within-group pairwise comparisons performed with Wilcoxon matched pairs test. For between-group comparisons, a Kruskal-Wallis ANOVA was used with Mann-Whitney U tests for pairwise comparisons. All statistical analyses were performed using Statistica (v13, TIBCO Software Inc., Palo Alto, CA) alpha set *a priori* to 0.05.

Prior to this study, there were no published data on LV E_es_ in a porcine model of spinal cord injury. However, in our group’s rodent studies of T2 SCI, we have reported a significant mean difference in E_es_ of 0.67 mmHg/μl with a pooled SD of 0.17 mmHg/μl between animals with T2 SCI and sham injury (i.e., control)^13^. For experiment 1, assuming a power of 0.95 and α of 0.05 we required a minimum of six animals per group to detect significant changes in E_es_ across four time-points (pre-SCI and every hour up to 4 hours post-SCI). We chose to include a minimum of seven animals per group to account for any discrepancies in placing the LV-PV catheter and spinal cord probes.

For experiment 2 examining the impacts of hemodynamic management, no published data in the porcine model of SCI had reported significant between-group differences in SCO_2_. However, our group had reported a pooled SD of SCO_2_ (expressed as % of baseline) of 40% in animals receiving vasopressor-based hemodynamic management^10^. With seven animals per group, we were powered to detect a difference of 29% between groups utilizing and SD of 40%, an α of 0.05 and power of 0.95.

## Data availability

The authors confirm that the data supporting the findings of this study are available within the paper and its Supplementary material

## Acknowledgments

We thank Dr. Robert Boushel for providing his equipment and expertise for the thermodilution method in these experiments.

## Sources of Funding

US Department of Defense (W81XWH017-1-0660), Craig Nielsen Foundation (459120), Michael Smith Foundation for Health Research (Trainee Award #17197)

## Author contributions

C.R.W. contributed to the conception of the study and its design, interpretation of data, drafting and final editing of the manuscript. A.M.W. contributed to the study design, data acquisition, analyses and interpretation, drafting and revision of the manuscript. N.M. contributed to the study design, data acquisition and revision of the manuscript. E.E. contributed to data acquisition and analyses, and revision of the manuscript. K.T., K.S., F.S., K.Sh. and K-T.K. contributed to the data acquisition and revision of the manuscript. B.K.K. contributed to the conception of the study and its design, interpretation of data, drafting and final editing of the manuscript.

## Competing interests

No conflicts of interest to report.

## Supplementary Materials

Supplemental Tables S1a-b, Tables S2a-e and Table S3

