## Supplemental Table for "Cardio-centric hemodynamic management improves spinal cord oxygenation and mitigates hemorrhage in acute spinal cord injury"

### Supplementary Materials

**Table S1a.** Experiment 1: Left ventricular (LV) load-dependent pressure-volume indices, and global hemodynamics

| | | **Pre-SCI** | **Post-SCI** | | | |
|---|---|---|---|---|---|---|
| | **RM-ANOVA** | ***Baseline*** | ***1 hr*** | ***2 hr*** | ***3 hr*** | ***4 hr*** |
| | | **LV volumetric measures** | | | | |
| **EDV** (ml) | **p=0.003** | 57.7 (9.9) | 56.5 (10.6) | 58.0 (11.4) | 60.6 (11.0) | 62.4 (13.7)** |
| **ESV** (ml) | **p=0.04** | 32.3 (6.9) | 30.2 (6.3) | 30.7 (6.4) | 31.1 (5.6) | 33.4 (7.2) |
| **SV** (ml) | p=0.10 | 27.4 (5.9) | 28.0 (5.9) | 29.0 (6.1) | 31.3 (6.4) | 30.7 (8.5) |
| **EF** (%) | p=0.27 | 49.5 (8.2) | 52.4 (6.8) | 52.8 (7.0) | 54.5 (7.5) | 52.3 (10.1) |
| | | **LV systolic pressure measures** | | | | |
| $\mathbf{P_{max}}$ (mmHg) | **p=0.003** | 101.0 (12.7) | 95.2 (11.0)** | 93.2 (10.4)*** | 93.4 (9.0)*** | 94.3 (10.2)** |
| $\mathbf{P_{es}}$ (mmHg) | **p<0.001** | 97.2 (15.9) | 88.0 (15.2)** | 85.5 (13.9)*** | 85.7 (12.7)*** | 87.1 (14.3)*** |
| $\mathbf{dp/dt_{max}}$ (mmHg/s) | p=0.24 | 1630 (194) | 1705 (156) | 1695 (179) | 1685 (168) | 1716 (215) |
| **Stroke work** (mmHgml) | p=0.24 | 2053 (387) | 2084 (561) | 2114 (560) | 2291 (578) | 2275 (721) |
| $\mathbf{E_a}$ (mmHg/ml) | **p=0.005** | 3.69 (1.09) | 3.17 (0.94) | 3.00 (0.90)* | 2.77 (0.81)** | 2.94 (2.94)** |
| | | **LV diastolic pressure measures** | | | | |
| $\mathbf{P_{ed}}$ (mmHg) | p=0.89 | 15.3 (3.7) | 14.6 (3.5) | 14.9 (3.5) | 14.6 (2.6) | 15.2 (3.2) |
| $\mathbf{dp/dt_{min}}$ (mmHg/s) | p=0.15 | -2357 (696) | -2223 (662) | -2182 (709) | -2150 (668) | -2212 (650) |
| **τ** (ms) | p=0.93 | 43 (7) | 42 (7) | 43 (8) | 42 (7) | 42 (6) |
| | | **Systemic and pulmonary hemodynamics** | | | | |
| **MAP** (mmHg) | **p<0.001** | 85.4 (13.2) | 79.5 (12.5)** | 77.4 (12.6)*** | 78.6 (11.2)*** | 78.9 (11.4)*** |
| **SBP** (mmHg) | **p=0.012** | 116.2 (12.5) | 111.1 (15.1)** | 110.0 (15.2)*** | 112.3 (13.5)* | 112.7 (13.9)* |
| **DBP** (mmHg) | **p<0.001** | 69.9 (14.3) | 63.6 (12.1)** | 61.1 (12.3)*** | 61.7 (11.1)*** | 62.0 (11.4)*** |
| $\mathbf{Q_{td}}$ (ml/min) | p=0.07 | 2685 (473) | 2902 (463) | 2528 (1126) | 2528 (1080) | 2572 (1085) |
| **HR** (bpm) | p=0.45 | 89.3 (19.0) | 94.6 (15.1) | 92.9 (14.3) | 91.1 (15.0) | 92.4 (14.0) |
| **TPR** (mmHg/L/min) | **p=0.003** | 37.4 (10.6) | 31.2 (6.8)** | 29.8 (6.8)** | 28.9 (7.3)*** | 29.9 (9.5)** |
| **mPAP** (mmHg) | p=0.15 | 21.1 (4.1) | 22.6 (4.7) | 22.6 (4.3) | 22.4 (3.6) | 22.6 (3.7) |
| **PASP** (mmHg) | p=0.12 | 31.5 (4.3) | 32.8 (5.3) | 33.0 (5.0) | 33.0 (4.5) | 32.9 (4.8) |

Values are means (SD). Time effects shown for one-way repeated measures ANOVA (RM-ANOVA), pairwise comparisons versus pre-SCI (baseline) performed using Fisher's LSD. EDV: end-diastolic volume; ESV: end-systolic volume; SV: stroke volume; EF: ejection fraction; $P_{max}$: maximal systolic pressure; $P_{es}$: end-systolic pressure; $dp/dt_{max}$: maximal rate of pressure generation; $E_a$: arterial elastance; $P_{ed}$: end-diastolic pressure; $dp/dt_{min}$: maximum rate of pressure decay; **τ**: rate constant of pressure decay; MAP: mean arterial pressure; SBP: systolic blood pressure; DBP: diastolic blood pressure; $Q_{td}$: cardiac output assessed with thermodilution; HR: heart rate; TPR: total peripheral resistance; mPAP: mean pulmonary arterial pressure; PASP: pulmonary arterial systolic pressure. *p<0.05 vs pre-SCI; **p<0.01 vs pre-SCI; ***p<0.001 vs pre-SCI.

**Table S1b.** Experiment 1: Left ventricular (LV) load-independent function and vascular-ventricular coupling

| | RM-ANOVA | Pre-SCI *Baseline* | Post-SCI *1 hr* | *2 hr* | *3 hr* | *4 hr* |
|---|---|---|---|---|---|---|
| **$E_{es}$** (mmHg/ml) | **p<0.001** | 2.66 (0.76) | 2.16 (0.72)*** | 2.17 (0.73)*** | 2.08 (0.62)*** | 2.11 (0.70)*** |
| **$EDPVR_{B1}$** | p=0.24 | 0.018 (0.008) | 0.014 (0.008) | 0.017 (0.007) | 0.015 (0.006) | 0.015 (0.007) |
| **PRSW** (mmHg) | **p=0.013** | 57.2 (13.8) | 46.7 (12.4)** | 48.2 (13.1)* | 47.0 (9.5)*** | 46.8 (11.7)*** |
| **$dp/dt_{max}$-EDV** (mmHg/s/ml) | **p<0.001** | 29.3 (12.5) | 21.5 (12.2)** | 23.3 (13.5)** | 18.8 (9.9)*** | 17.7 (9.6)*** |
| **$E_a/E_{es}$** | p=0.58 | 1.42 (0.39) | 1.52 (0.41) | 1.43 (0.37) | 1.36 (0.35) | 1.43 (0.44) |

Values are means (SD). Statistics are identical to those in Table S1a. $E_{es}$: end-systolic elastance, index of contractility; $EDPVR_{B1}$: end-diastolic pressure-volume relationship slope; PRSW: preload-recruitable stroke work; $dp/dt_{max}$-EDV: maximal rate of pressure generation for a given end-diastolic volume; $E_a/E_{es}$: vascular-ventricular coupling, arterial elastance -to- end-systolic elastance. *p<0.05 vs pre-SCI; **p<0.01 vs pre-SCI; ***p<0.001 vs pre-SCI.

**Table S2a.** Experiment 2: Left ventricular (LV) load-independent function

| | RM-ANOVA EFFECTS | | | GROUP | PRE-SCI | POST-SCI | | | | |
|---|---|---|---|---|---|---|---|---|---|---|
| | *time* | *group* | *gr × time* | | *Baseline* | *30 min (pre-drug)* | *1 hr* | *2 hr* | *3 hr* | *4 hr* |
| **$E_{es}$** | p=0.098 | **p<0.001** | **p<0.001** | CON | 2.66 (0.76) | - | 2.16 (0.72) | 2.17 (0.73) | 2.08 (0.62) | 2.11 (0.70) |
| (mmHg/ml) | | | | DOB+ † | 1.93 (0.63) | 1.90 (0.45) | 3.13 (0.90)* | 3.26 (1.26)* | 3.54 (1.48)* | 4.20 (1.96)** |
| | | | | DOB- † | 3.36 (1.64) | 2.15 (0.30) | 3.61 (1.02)** | 3.24 (0.14)* | 3.05 (0.19) | 3.34 (0.16)* |
| | | | | NE †† | 2.26 (0.94) | 2.21 (0.61) | 4.39 (1.70)** [a] | 4.99 (2.35)*** [a] | 4.83 (2.27)** [a] | 4.55 (1.92)*** [a] |
| **$EDPVR_{B1}$** | p=0.17 | p=0.80 | p=0.56 | CON | 0.012 (0.009) | - | 0.021 (0.014) | 0.020 (0.014) | 0.020 (0.017) | 0.020 (0.018) |
| | | | | DOB+ | 0.019 (0.009) | 0.026 (0.019) | 0.023 (0.011) | 0.021 (0.009) | 0.020 (0.007) | 0.020 (0.009) |
| | | | | DOB- | 0.027 (0.017) | 0.031 (0.011) | 0.034 (0.017) | 0.030 (0.011) | 0.033 (0.014) | 0.023 (0.005) |
| | | | | NE | 0.022 (0.008) | 0.023 (0.012) | 0.028 (0.011) | 0.027 (0.012) | 0.024 (0.009) | 0.025 (0.008) |
| **PRSW** | p=0.055 | **p<0.001** | **p<0.001** | CON | 29.3 (12.6) | - | 21.5 (12.2) | 23.3 (13.5) | 18.8 (9.9) | 17.7 (9.6) |
| (mmHg) | | | | DOB+ ††† | 15.4 (8.6) | 20.4 (6.6) | 49.5 (14.8)*** | 50.5 (26.9)*** [a] | 58.5 (4.0)*** [a] | 59.4 (10.7)*** [a] |
| | | | | DOB- †† | 36.6 (7.2) | 36.0 (5.6) | 49.2 (7.0)** | 50.4 (5.3)** | 43.4 (6.7)** | 45.3 (7.6)** |
| | | | | NE ††† | 25.4 (13.1) | 24.8 (20.3) | 52.9 (28.3)*** [a] | 77.2 (54.8)*** [a] | 101.6(120.9)*** [a] | 99.9 (99.5)*** [a] |
| **$dp/dt_{max}$-EDV** | p=0.097 | **p<0.001** | **p<0.001** | CON | 57.2 (13.8) | - | 46.7 (12.4) | 48.2 (13.1) | 47.0 (9.5) | 46.8 (11.7) |
| (mmHg/s/ml) | | | | DOB+ † | 38.4 (15.3) | 38.5 (11.9) | 75.8 (5.5)** | 81.2 (4.8)** | 83.9 (7.5)** [a] | 83.3 (11.5)** [a] |
| | | | | DOB- ††† | 59.9 (20.0) | 48.4 (7.6) | 66.3 (15.8)*** | 68.6 (8.1)*** | 66.7 (14.9)** | 66.6 (15.6)*** |
| | | | | NE ††† | 48.8 (18.7) | 46.7 (21.4) | 78.9 (24.7)** [a] | 78.5 (20.2)*** | 84.9 (28.5)*** [a] | 88.2 (26.3)*** [a] |
| **$E_a/E_{es}$** | p=0.41 | **p<0.001** | **p<0.001** | CON | 1.42 (0.39) | - | 1.52 (0.41) | 1.43 (0.37) | 1.36 (0.35) | 1.43 (0.44) |
| | | | | DOB+ †† | 1.91 (0.57) | 1.62 (0.51) | 0.89 (0.30)*** | 0.91 (0.34)*** [a] | 0.87 (0.29)*** | 0.78 (0.45)*** [a] |
| | | | | DOB- †† | 1.27 (0.58) | 1.36 (0.15) | 0.85 (0.20)*** | 0.88 (0.06)** [a] | 0.98 (0.12)** | 0.89 (0.04)** |
| | | | | NE †† | 1.74 (0.87) | 1.57 (0.59) | 0.96 (0.69)** | 0.96 (0.79)** | 0.95 (0.72)*** | 0.94 (0.76)*** |

Values are means (SD). CON: control (*n*=8); DOB+: high-dose (i.e. ≥2.5 µg/kg/min; *n*=4) Dobutamine; DOB-: low-dose Dobutamine (i.e. ≤0.5 µg/kg/min; *n*=3); NE: Norepinephrine (*n*=7). Effects from 2-way RM-ANOVA are shown for time, group and interaction (group x time). Significant effects for 1-way repeated measures ANOVA are shown for each experimental group (in "GROUP" column) with symbols. Tukey's HSD was performed post-hoc for between-group comparisons, and Fisher's LSD for within-group comparisons (i.e. versus 30 mins / pre-drug). See Tables S1a and S1b for additional abbreviations. Between-group comparisons: [a] p<0.05 vs CON; [b] p<0.05 vs DOB - ; [c] p<0.05 vs NE. Within-group comparison to 30 mins (i.e. pre-drug): *p<0.05; **p<0.01; ***p<0.001. Within-group effect of time from 30 mins to 4 hrs post-SCI: †p<0.05; ††p<0.01; †††p<0.001.

**Table S2b.** Experiment 2: Left ventricular (LV) load-dependent pressure-volume indices for systolic and diastolic function

| | RM-ANOVA EFFECTS | | | GROUP | PRE-SCI | POST-SCI | | | | |
|---|---|---|---|---|---|---|---|---|---|---|
| | *time* | *group* | *gr × time* | | *Baseline* | *30 min (pre-drug)* | *1 hr* | *2 hr* | *3 hr* | *4 hr* |
| **LV volumetric measures** | | | | | | | | | | |
| **EDV** | p=0.62 | **p=0.006** | **p=0.003** | CON ††† | 57.7 (9.9) | 54.1 (8.6) | 56.5 (10.6) | 58.0 (11.4)* | 60.6 (11.0)*** | 62.4 (13.7)*** |
| (ml) | | | | DOB+ †† | 63.8 (8.2) | 65.3 (7.5) | 58.2 (7.1)** | 58.0 (6.1)** | 60.4 (4.6)* | 61.6 (5.2)* |
| | | | | DOB- | 52.8 (3.3) | 52.5 (2.0) | 48.6 (3.8) | 51.6 (2.7) | 54.9 (4.6) | 56.6 (5.2) |
| | | | | NE | 61.4 (10.3) | 60.7 (12.1) | 54.9 (9.6) | 50.1 (11.1) | 53.4 (12.1) | 54.8 (12.9) |
| **ESV** | p=0.57 | **p<0.001** | **p<0.001** | CON †† | 32.3 (6.9) | 28.6 (4.8) | 30.2 (6.3) | 30.7 (6.4)* | 31.1 (5.6)* | 33.4 (7.2)*** |
| (ml) | | | | DOB+ ††† | 36.8 (9.3) | 36.1 (8.1) | 22.7 (5.3)*** | 21.9 (6.8)*** | 22.9 (5.4)*** | 22.6 (4.7)*** |
| | | | | DOB- † | 30.8 (5.5) | 28.3 (5.5) | 22.9 (6.6)** | 24.7 (6.6)* | 27.5 (7.9) | 28.3 (8.3) |
| | | | | NE † | 36.7 (12.0) | 34.8 (12.5) | 26.0 (8.7)* | 23.6 (8.7)** | 25.6 (10.7)** | 26.1 (11.4)* |
| **SV** | p=0.58 | **p<0.001** | p=0.11 | CON | 27.4 (5.9) | 27.8 (5.7) | 28.0 (5.9) | 29.0 (6.1) | 31.3 (6.4) | 30.7 (8.5) |
| (ml) | | | | DOB+ ††† | 28.7 (1.7) | 31.4 (1.8) | 37.2 (2.4)***[a] | 37.6 (2.8)***[c] | 38.8 (1.2)***[c] | 40.3 (2.7)*** |
| | | | | DOB- † | 24.6 (2.3) | 27.4 (2.6) | 29.6 (3.9) | 30.1 (2.5)* | 30.8 (2.8)** | 31.9 (3.0)** |
| | | | | NE | 27.6 (3.8) | 28.5 (28.5) | 30.6 (4.4) | 28.0 (5.2) | 29.8 (4.4) | 30.9 (4.7) |
| **EF** | p=0.23 | **p<0.001** | **p<0.001** | CON | 49.5 (8.2) | 52.1 (6.1) | 52.4 (6.8) | 52.8 (7.0) | 54.5 (7.5) | 52.3 (10.1) |
| (%) | | | | DOB+ ††† | 46.5 (7.2) | 49.4 (6.6) | 65.2 (4.3)*** | 66.4 (8.1)*** | 65.7 (5.8)*** | 66.9 (5.0)*** |
| | | | | DOB- †† | 46.0 (6.9) | 51.2 (6.5) | 59.2 (7.6)*** | 57.0 (7.5)*** | 55.4 (8.3)** | 55.6 (8.4)** |
| | | | | NE †† | 49.1 (9.1) | 53.1 (7.6) | 60.2 (11.7)*** | 59.5 (10.0)** | 59.1 (10.8)** | 59.4 (10.7)** |
| **LV systolic pressure measures** | | | | | | | | | | |
| **$P_{max}$** | **p=0.01** | **p<0.001** | **p<0.001** | CON | 101.0 (12.7) | 83.0 (11.3) | 88.0 (15.2) | 85.5 (13.9) | 85.7 (12.7) | 87.1 (14.3) |
| (mmHg) | | | | DOB+ ††† | 103.7 (8.0) | 90.9 (6.8) | 96.6 (11.9)** | 99.2 (11.7)*** [a] | 106.9 (10.9)*** [ab] | 106.3 (15.3)*** [a] |
| | | | | DOB- † | 95.1 (4.4) | 79.4 (11.7) | 86.3 (4.8)* | 85.7 (4.5) | 90.9 (1.9)* | 94.6 (4.1)** |
| | | | | NE ††† | 97.1 (12.8) | 84.3 (16.9) | 101.1 (11.2)*** [a] | 97.5 (15.2)*** [a] | 101.5 (16.6)*** [a] | 99.1 (18.0)** |
| **$P_{es}$** | p=0.22 | **p<0.001** | **p=0.008** | CON | 97.2 (15.9) | 91.5 (8.6) | 95.2 (11.0) | 93.2 (10.4) | 93.4 (9.0) | 94.3 (10.2) |
| (mmHg) | | | | DOB+ † | 97.9 (7.9) | 99.0 (4.2) | 108.3 (8.3) | 111.1 (7.4) | 116.3 (7.4)** | 116.6 (11.7)** |
| | | | | DOB- | 89.4 (6.6) | 88.1 (8.3) | 95.1 (5.6) | 93.8 (5.0) | 96.0 (2.8) | 98.7 (4.1) |
| | | | | NE ††† | 91.8 (14.2) | 92.7 (14.2) | 109.5 (7.8)*** | 108.1 (8.4)*** | 110.6 (8.5)*** | 108.3 (9.5)*** |
| **$dp/dt_{max}$** | **p=0.004** | **p<0.001** | **p<0.001** | CON | 1630 (194) | 1731 (235) | 1705 (156) | 1695 (179) | 1685 (168) | 1716 (215) |
| (mmHg/s) | | | | DOB+ ††† | 1696 (354) | 1683 (345) | 3168 (726)*** [a] | 3523 (615)*** [a] | 3722 (241)*** | 3652 (373)*** [a] |
| | | | | DOB- †† | 1739 (111) | 1729 (235) | 2540 (551)*** | 2278 (504)** | 2178 (511)* | 2337 (621)** |
| | | | | NE †† | 1675 (238) | 1667 (195) | 3042 (752)** [a] | 3508 (1316)*** [a] | 3693 (2081)*** [a] | 3551 (1769)*** [a] |

**Table S2b *continued.*** Experiment 2: Left ventricular (LV) load-dependent pressure-volume indices for systolic and diastolic function

| | RM-ANOVA EFFECTS | | | GROUP | PRE-SCI | POST-SCI | | | | |
|---|---|---|---|---|---|---|---|---|---|---|
| | *time* | *group* | *gr × time* | | *Baseline* | *30 min (pre-drug)* | *1 hr* | *2 hr* | *3 hr* | *4 hr* |
| **Stroke work** | **p=0.002** | **p<0.001** | **p=0.004** | CON | 2053 (387) | 1969 (520) | 2084 (561) | 2114 (560) | 2291 (578) | 2275 (721) |
| (mmHg*ml) | | | | DOB+ ††† | 2351 (189) | 2428 (107) | 3287 (280)*** [a] | 3401 (99)*** [abc] | 3649 (86)*** [abc] | 3751 (166)*** [abc] |
| | | | | DOB- † | 1954 (46) | 1963 (288) | 2335 (317)* | 2340 (219)* | 2413 (268)** | 2547 (354)** |
| | | | | NE †† | 2078 (307) | 2111 (271) | 2706 (618)** | 2464 (433)* | 2642 (351)** | 2695 (368)** |
| **$E_a$** | p=0.73 | p=0.95 | **p=0.029** | CON | 3.69 (1.09) | 3.11 (0.81) | 3.17 (0.94) | 3.00 (0.90) | 2.77 (0.81) | 2.94 (2.94) |
| (mmHg/ml) | | | | DOB+ | 3.43 (0.42) | 2.91 (0.28) | 2.62 (0.51) | 2.66 (0.45) | 2.76 (0.32) | 2.67 (0.56) |
| | | | | DOB- | 3.67 (0.54) | 2.89 (0.23) | 2.93 (0.21) | 2.86 (0.10) | 2.97 (0.24) | 2.97 (0.18) |
| | | | | NE †† | 3.23 (0.72) | 2.84 (0.99) | 3.29 (0.80)** | 3.45 (0.93)*** | 3.36 (0.74)** | 3.20 (0.84)* |
| **LV diastolic pressure measures** | | | | | | | | | | |
| **$P_{ed}$** | p=0.25 | **p=0.001** | **p=0.028** | CON | 15.3 (3.7) | 13.3 (2.7) | 14.6 (3.5) | 14.9 (3.5) | 14.6 (2.6) | 15.2 (3.2) |
| (mmHg) | | | | DOB+ | 14.0 (4.3) | 15.0 (4.1) | 13.5 (5.9) | 13.4 (3.5) | 14.9 (5.1) | 15.0 (5.3) |
| | | | | DOB- † | 10.4 (4.2) | 10.1 (4.0) | 8.8 (5.6) | 9.6 (4.1) | 11.5 (3.0) | 13.4 (2.3)* |
| | | | | NE | 13.8 (2.8) | 13.4 (3.0) | 15.0 (4.9) | 13.6 (4.9) | 14.6 (5.3) | 14.5 (5.3) |
| **$dp/dt_{min}$** | **p=0.036** | **p<0.001** | **p=0.009** | CON † | -2357 (696) | -2045 (596) | -2223 (662)** | -2182 (709)* | -2150 (668)* | -2212 (650)** |
| (mmHg/s) | | | | DOB+ †† | -3152 (706) | -2597 (425) | -2988 (533) | -3311 (654)* | -3596 (760)** | -3482 (1022)** |
| | | | | DOB- † | -3433 (155) | -2775 (455) | -3426 (136)* | -3434 (160)* | -3328 (258)* | -3435 (139)* |
| | | | | NE † | -3277 (1073) | -2789 (911) | -3155 (864)* | -3121 (975) | -3406 (988)*** | -3303 (1015)** |
| **τ** | p=0.07 | **p=0.046** | **p=0.026** | CON | 43 (7) | 42 (6) | 42 (7) | 43 (8) | 42 (7) | 42 (6) |
| (ms) | | | | DOB+ † | 34 (5) | 36 (6) | 30 (4)** | 30 (2)** | 31 (2)** | 32 (3)* |
| | | | | DOB- ††† | 32 (5) | 33 (4) | 31 (6)* | 32 (6) | 36 (6)* | 36 (5)** |
| | | | | NE | 39 (5) | 41 (7) | 41 (10) | 40 (11) | 39 (10) | 42 (8) |

Values are means (SD). CON: control (*n*=8); DOB+: high-dose (i.e. ≥2.5 µg/kg/min; *n*=4) Dobutamine; DOB-: low-dose Dobutamine (i.e. ≤0.5 µg/kg/min; *n*=3); NE: Norepinephrine (*n*=7). Statistics are identical to those outlined in Table S2a. See Supplemental Tables S1a and S1b for additional abbreviations. Between-group comparisons: [a] p<0.05 vs CON; [b] p<0.05 vs DOB - ; [c] p<0.05 vs NE. Within-group comparison to 30 mins (i.e. pre-drug): *p<0.05; **p<0.01; ***p<0.001. Within-group effect of time from 30 mins to 4 hrs post-SCI: †p<0.05; ††p<0.01; †††p<0.001.

**Table S2c.** Experiment 2: Global and pulmonary hemodynamics

| | RM-ANOVA EFFECTS | | | GROUP | PRE-SCI | POST-SCI | | | | |
|---|---|---|---|---|---|---|---|---|---|---|
| | *time* | *group* | *gr × time* | | *Baseline* | *30 min* | *1 hr* | *2 hr* | *3 hr* | *4 hr* |
| **Systemic blood pressure and vascular resistance measures** | | | | | | | | | | |
| **MAP** | p=0.07 | **p<0.001** | **p<0.001** | CON | 85.4 (13.2) | 77.1 (11.4) | 79.5 (12.5) | 77.4 (12.6) | 78.6 (11.2) | 78.9 (11.4) |
| (mmHg) | | | | DOB+ ††† | 88.1 (7.8) | 82.6 (2.7) | 91.5 (3.8)** | 94.7 (5.0)*** **a** | 100.3 (1.9)*** **a** | 99.1 (6.2)*** **a** |
| | | | | DOB- † | 80.4 (5.5) | 75.8 (2.2) | 79.7 (2.4) | 79.2 (2.1) | 81.3 (3.0)* | 82.4 (3.0)** |
| | | | | NE ††† | 76.3 (11.7) | 72.7 (12.9) | 88.2 (5.6)*** | 86.8 (9.0)*** | 88.2 (11.3)*** | 85.7 (12.7)*** |
| **SBP** | **p=0.006** | **p<0.001** | **p<0.001** | CON † | 116.2 (12.5) | 108.1 (13.2) | 111.1 (15.1)* | 110.0 (15.2) | 112.3 (13.5)** | 112.7 (13.9)** |
| (mmHg) | | | | DOB+ †† | 126.9 (13.4) | 123.6 (5.5) | 141.3 (10.7)** **a** | 147.9 (7.1)** **ab** | 154.2 (10.7)*** **abc** | 150.5 (19.4)*** **a** |
| | | | | DOB- † | 112.8 (2.9) | 108.3 (8.2) | 113.7 (12.4)* | 112.2 (8.8) | 113.3 (8.8)* | 116.5 (7.4)** |
| | | | | NE ††† | 108.0 (16.1) | 104.9 (16.9) | 128.1 (11.5)*** | 125.3 (16.2)*** | 126.3 (17.3)*** | 123.4 (18.9)*** |
| **DBP** | p=0.69 | **p<0.001** | **p=0.037** | CON | 69.9 (14.3) | 61.6 (11.4) | 63.6 (12.1) | 61.1 (12.3) | 61.7 (11.1) | 62.0 (11.4) |
| (mmHg) | | | | DOB+ †† | 68.7 (6.6) | 62.0 (2.4) | 66.5 (2.8) | 68.1 (5.2)* | 73.3 (5.9)** | 73.4 (7.3)** |
| | | | | DOB- | 64.3 (9.1) | 59.6 (4.2) | 62.7 (5.8) | 62.7 (5.1) | 65.3 (8.9) | 65.4 (7.2) |
| | | | | NE † | 60.6 (11.1) | 56.6 (11.8) | 68.2 (8.7)** | 67.6 (8.8)** | 69.1 (13.0)** | 66.9 (14.9)* |
| **TPR** | p=0.94 | p=0.76 | p=0.15 | CON | 37.4 (10.6) | 29.3 (6.5) | 31.2 (6.8) | 29.8 (6.8) | 28.9 (7.3) | 29.9 (9.5) |
| (mmHg/l/min) | | | | DOB+ | 34.0 (8.3) | 29.0 (4.7) | 25.5 (7.5) | 26.5 (4.5) | 28.1 (4.7) | 26.6 (6.3) |
| | | | | DOB- | 35.9 (0.9) | 31.4 (6.2) | 26.4 (7.6) | 27.4 (5.3) | 31.6 (6.9) | 31.4 (4.4) |
| | | | | NE | 29.4 (4.2) | 27.1 (7.6) | 30.5 (9.0) | 30.3 (12.0) | 29.8 (9.4) | 28.8 (10.0) |
| **Cardiac output and heart rate measures** | | | | | | | | | | |
| **$Q_{pv}$** | p=0.23 | **p=0.002** | p=0.22 | CON | 2386 (486) | 2691 (482) | 2601 (491) | 2665 (549) | 2823 (633) | 2831 (807) |
| (ml/min) | | | | DOB+ † | 2684 (577) | 2897 (465) | 3759 (822)* | 3636 (480)* | 3640 (545)* | 3847 (711)** |
| | | | | DOB- † | 2241 (131) | 2479 (473) | 3179 (835)** | 2960 (565)* | 2652 (559) | 2664 (454) |
| | | | | NE | 2637 (563) | 2780 (636) | 3156 (1127) | 3193 (1037) | 3192 (910) | 3280 (1027) |
| **HR** | p=0.61 | p=0.36 | p=0.13 | CON | 89.3 (19.0) | 98.8 (17.4) | 94.6 (15.1) | 92.9 (14.3) | 91.1 (15.0) | 92.4 (14.0) |
| (bpm) | | | | DOB+ | 91.1 (14.6) | 91.7 (13.2) | 101.2 (17.4) | 96.2 (6.2) | 94.7 (10.6) | 101.6 (24.0) |
| | | | | DOB- | 91.9 (14.2) | 91.6 (11.7) | 101.9 (20.9) | 97.1 (13.9) | 86.3 (11.8) | 82.2 (6.7) |
| | | | | NE | 95.5 (14.8) | 96.6 (13.4) | 101.7 (25.5) | 113.7 (31.9) | 108.3 (34.8) | 107.2 (35.0) |

**Table S2c *continued.*** Experiment 2: Global and pulmonary hemodynamics

| | RM-ANOVA EFFECTS | | | GROUP | PRE-SCI | POST-SCI | | | | |
|---|---|---|---|---|---|---|---|---|---|---|
| | *time* | *group* | *gr × time* | | *Baseline* | *30 min* | *1 hr* | *2 hr* | *3 hr* | *4 hr* |
| **Pulmonary pressure measures** | | | | | | | | | | |
| **mPAP** | **p=0.017** | **p=0.028** | **p=0.018** | CON | 21.1 (4.1) | 22.0 (4.6) | 22.6 (4.7) | 22.6 (4.3) | 22.4 (3.6) | 22.6 (3.7) |
| (mmHg) | | | | DOB+ | 15.4 (2.4) | 14.7 (2.2)[a] | 14.7 (2.9)[a] | 14.9 (2.7) | 15.3 (1.8)[a] | 15.7 (2.5)[a] |
| | | | | DOB- | 16.7 (2.1) | 15.8 (1.4) | 16.1 (1.3) | 15.8 (1.3) | 16.0 (1.5) | 16.4 (1.6) |
| | | | | NE ††† | 16.5 (3.5) | 16.3 (3.3)[a] | 18.6 (4.8)* | 20.9 (4.9)*** | 19.2 (5.2)** | 18.5 (4.8)* |
| **PASP** (mmHg) | p=0.45 | **p<0.001** | **p=0.017** | CON | 31.5 (4.3) | 31.6 (5.0) | 32.8 (5.3) | 33.0 (5.0) | 33.0 (4.5) | 32.9 (4.8) |
| | | | | DOB+ | 27.9 (4.9) | 28.2 (4.6) | 28.2 (4.6) | 30.2 (5.6) | 30.0 (5.7) | 29.6 (6.0) |
| | | | | DOB- | 27.8 (3.2) | 27.5 (3.1) | 27.5 (3.1) | 27.7 (2.8) | 27.6 (2.9) | 27.6 (3.1) |
| | | | | NE †† | 25.2 (3.7) | 25.3 (4.0) | 29.1 (6.9)** | 31.5 (7.4)*** | 30.0 (7.6)** | 29.3 (6.7)** |

Values are means (SD). CON: control (*n*=8); DOB+: high-dose (i.e. ≥2.5 µg/kg/min; *n*=4) Dobutamine; DOB-: low-dose Dobutamine (i.e. ≤0.5 µg/kg/min; *n*=3); NE: Norepinephrine (*n*=7). Statistics are identical to those outlined in Table S2a. $Q_{pv}$: cardiac output assessed with LV pressure-volume catheterization. See Tables S1a and S1b for additional abbreviations. Between-group comparisons: [a] p<0.05 vs CON; [b] p<0.05 vs DOB - ; [c] p<0.05 vs NE. Within-group comparison to 30 mins (i.e. pre-drug): *p<0.05; **p<0.01; ***p<0.001. Within-group effect of time: †p<0.05; ††p<0.01; †††p<0.001.

**Table S2d.** Experiment 2: Intraparenchymal spinal cord oxygenation, blood flow, pressure and metabolism from 1.2 cm probes and 3.2 cm probes

| | RM-ANOVA EFFECTS | | | GROUP | PRE-SCI | POST-SCI | | | | |
|---|---|---|---|---|---|---|---|---|---|---|
| | *time* | *group* | *gr × time* | | *Baseline* | *30 min (pre-drug)* | *1 hr* | *2 hr* | *3 hr* | *4 hr* |
| **Spinal cord oxygenation, blood flow and cord pressure at 1.2 cm** | | | | | | | | | | |
| **$SCO_2$ 1.2cm** | p=0.058 | **p<0.001** | **p<0.001** | CON ††† | 21.9 (13.4) | 0.6 (0.1) | 0.6 (0.1) | 0.6 (0.1) | 18.9 (13.4)*** | 21.9 (13.3)*** |
| (mmHg) | | | | DOB+ ††† | 22.9 (6.7) | 1.7 (1.7) | 7.4 (13.2) | 10.2 (18.2) | 42.5 (10.2)*** | 47.5 (9.9)*** |
| | | | | DOB- | 25.5 (1.2) | 1.5 (1.5) | 4.8 (7.3) | 6.8 (10.7) | 17.8 (14.3) | 20.8 (11.1) |
| | | | | NE †† | 27.0 (13.0) | 0.6 (0.1) | 0.9 (0.5) | 2.3 (4.1) | 26.2 (24.5)* | 26.4 (25.3)* |
| **$SCO_2$ 1.2cm** | p=0.31 | **p<0.001** | **p=0.026** | CON ††† | - | -95 (4) | -96 (4) | -96 (3) | -35 (51)** | -8 (63)*** |
| (%Δ from | | | | DOB+ ††† | - | -94 (5) | -97 (0) | -96 (1) | 89 (27)*** [ab] | 112 (27)*** [abc] |
| pre-SCI) | | | | DOB- | - | -94 (7) | -81 (30) | -73 (44) | -54 (16) | -42 (3)* |
| | | | | NE ††† | - | -97 (1) | -95 (5) | -80 (29) | -6 (72)*** | -6 (74)*** |
| **SCBF 1.2cm** | **p=0.008** | **p<0.001** | p=0.56 | CON ††† | 178 (66) | 66 (23) | 86 (47) | 70 (30) | 255 (45)*** | 299 (112)*** |
| (a.u.) | | | | DOB+ ††† | 356 (115) | 191 (94) | 261 (164)** | 328 (275)*** [a] | 327 (98)*** | 364 (119)*** |
| | | | | DOB- | 254 (164) | 135 (138) | 165 (182) | 134 (147) | 386 (341) | 329 (262) |
| | | | | NE †† | 248 (65) | 80 (54) | 120 (76) | 111 (81) | 300 (204)*** | 342 (247)*** |
| **SCBF 1.2cm** | p=0.125 | **p<0.001** | **p=0.003** | CON ††† | - | -52 (42) | -51 (37) | -58 (28) | 31 (42)*** | 35 (35)*** |
| (%Δ from | | | | DOB+ | - | -47 (19) | -31 (27) | -26 (35) | -14 (37) | -3 (48) |
| pre-SCI) | | | | DOB- | - | -80 (9) | -78 (10) | -82 (11) | 23 (77) | 6 (53) |
| | | | | NE † | - | -62 (32) | -47 (37) | -63 (21) | -12 (44) *** | -1 (54) ** |
| **SCP 1.2cm** | p=0.27 | **p<0.001** | p=0.20 | CON ††† | 9.5 (3.7) | 33.1 (19.8) | 32.3 (15.5) | 31.7 (11.5) | 11.6 (5.4)*** | 11.5 (5.0)*** |
| (mmHg) | | | | DOB+ | 6.4 (3.7) | 14.7 (6.0) | 17.8 (9.7) | 24.2 (19.9) | 8.9 (1.5) | 9.5 (1.9) |
| | | | | DOB- | 6.4 (4.3) | 20.1 (15.9) | 20.0 (17.1) | 16.0 (17.5) | 9.2 (7.2) | 10.6 (5.9) |
| | | | | NE ††† | 8.0 (5.0) | 20.2 (8.8) | 22.4 (8.4) | 26.1 (9.9) | 11.1 (3.4) ** | 10.9 (3.0) ** |
| **SCP 1.2cm** | p=0.79 | **p<0.001** | p=0.59 | CON †† | - | 345 (372) | 338 (351) | 348 (378) | 20 (28)** | 20 (21)** |
| (%Δ from | | | | DOB+ † | - | 236 (302) | 306 (384) | 399 (434) | 98 (154) | 133 (227) |
| pre-SCI) | | | | DOB- | - | 173 (186) | 152 (222) | 73 (176) | 46 (33) | 80 (30) |
| | | | | NE †† | - | 188 (185) | 213 (208) | 272 (283) | 39 (75) * | 32 (47) * |

**Table S2d *continued.*** Experiment 2: Intraparenchymal spinal cord oxygenation, blood flow, pressure and metabolism from 1.2 cm probes and 3.2 cm probes

| | RM-ANOVA EFFECTS | | | GROUP | PRE-SCI | POST-SCI | | | | |
|---|---|---|---|---|---|---|---|---|---|---|
| | *time* | *group* | *gr × time* | | *Baseline* | *30 min (pre-drug)* | *1 hr* | *2 hr* | *3 hr* | *4 hr* |
| **Spinal cord oxygenation, blood flow and cord pressure at 3.2 cm** | | | | | | | | | | |
| **$SCO_2$ 3.2cm** | **p<0.001** | **p<0.001** | p=0.22 | CON ††† | 25.3 (10.9) | 10.3 (9.2) | 17.9 (16.4) | 23.4 (14.4) | 34.7 (13.9)*** | 42.2 (11.3) *** |
| (mmHg) | | | | DOB+ ††† | 35.4 (5.5) | 14.4 (5.4) | 30.2 (1.3) | 55.5 (23.6) [a] | 57.2 (11.0)*** | 57.7 (8.3) *** |
| | | | | DOB- | 22.5 (14.0) | 15.4 (5.7) | 16.7 (2.7) | 19.5 (2.1) | 23.1 (16.9) [b] | 31.6 (17.7) |
| | | | | NE ††† | 39.7 (20.0) | 32.3 (12.6) [a] | 44.2 (19.9) [a] | 49.9 (17.6) [a] | 54.8 (17.2)*** [c] | 57.5 (16.3)*** |
| **$SCO_2$ 3.2cm** | p=0.21 | **p<0.001** | p=0.51 | CON †† | - | -47 (42) | -37 (45) | 2 (65) | 27 (59)* | 62 (71)*** |
| (%Δ from | | | | DOB+ † | - | -42 (36) | -8 (16) | 22 (18) | 35 (40)** | 40 (50)** |
| pre-SCI) | | | | DOB- | - | -8 (56) | -42 (5) [b] | -41 (3) | -46 (58) | -23 (51) |
| | | | | NE | - | -29 (27) | -4 (40) | 13 (42) | 22 (32)** | 28 (32)*** |
| **SCBF 3.2cm** | p=0.18 | p=0.57 | p=0.92 | CON † | 180 (65) | 214 (110) | 191 (82) | 201 (67) | 259 (51) | 294 (69)* |
| (a.u.) | | | | DOB+ †† | 651 (523) | 373 (149) | 415 (156) | 490 (157) | 631 (223)** | 660 (232)** |
| | | | | DOB- | 222 (102) | 330 (178) | 348 (194) | 326 (181) | 285 (151) | 340 (191) |
| | | | | NE † | 390 (267) | 958 (1059) | 1159 (1381) | 1312 (1647) | 1152 (1312) | 1153 (1064) |
| **SCBF 3.2cm** | p=0.38 | p=0.23 | p=0.50 | CON †† | - | 18 (43) | 11 (38) | 22 (43) | 57 (45)* | 81 (60)** |
| (%Δ from | | | | DOB+ † | - | 16 (44) | 30 (52) | 58 (68) | 100 (93)** | 96 (105)** |
| pre-SCI) | | | | DOB- | - | 42 (27) | 51 (43) | 42 (41) | 25 (30) | 51 (49) |
| | | | | NE | - | 149 (271) | 203 (347) | 243 (409) | 189 (311) | 183 (240) |
| **SCP 3.2cm** | p=0.92 | p=0.67 | p=0.41 | CON ††† | 14.6 (6.0) | 12.6 (5.8) | 12.0 (5.6) | 12.0 (6.2) | 13.4 (5.2) | 14.1 (5.4)** |
| (mmHg) | | | | DOB+ | 13.0 (0.9) | 14.3 (2.1) | 13.2 (2.3) | 10.4 (6.4) | 10.3 (5.4) | 11.3 (4.3) |
| | | | | DOB- | 11.2 (0.4) | 12.3 (0.78) | 11.7 (1.7) | 11.5 (1.9) | 15.6 (5.7) | 15.2 (4.3) |
| | | | | NE | 11.0 (4.6) | 15.4 (11.1) | 14.7 (10.7) | 14.9 (12.7) | 13.2 (4.1) | 13.2 (2.7) |
| **SCP 3.2cm** | p=0.30 | p=0.89 | p=0.75 | CON †† | - | -16 (17) | -20 (15) | -22 (17) | -8 (13)** | -2 (16)** |
| (%Δ from | | | | DOB+ | - | 10 (15) | 2 (16) | -20 (49) | -22 (39) | -14 (30) |
| pre-SCI) | | | | DOB- | - | 10 (7) | 4 (16) | 3 (18) | 40 (56) | 36 (43) |
| | | | | NE | - | 51 (131) | 47 (129) | 53 (154) | 34 (59) | 34 (26) |

Values are means (SD). CON: control (*n*=8); DOB+: high-dose (i.e. ≥2.5 μg/kg/min; *n*=4) Dobutamine; DOB-: low-dose Dobutamine (i.e. ≤0.5 μg/kg/min; *n*=3); NE: Norepinephrine (*n*=7). Statistics are identical to those outlined in Table S2a. $SCO_2$: spinal cord oxygenation; SCBF: spinal cord blood flow; SCP: spinal cord pressure; a.u.: arbitrary units. Between-group comparisons: [a] p<0.05 vs CON; [b] p<0.05 vs DOB - ; [c] p<0.05 vs NE. Within-group comparison to 30 mins (i.e. pre-drug): *p<0.05; **p<0.01; ***p<0.001. Within-group effect of time from 30 mins to 4 hrs post-SCI: †p<0.05; ††p<0.01; †††p<0.001.

**Table S2e.** Experiment 2: Microdialysis measures of spinal cord metabolism

| | RM-ANOVA EFFECTS | | | GROUP | PRE-SCI | POST-SCI | | | | |
|---|---|---|---|---|---|---|---|---|---|---|
| | *time* | *group* | *gr × time* | | *Baseline* | *30 min (pre-drug)* | *1 hr* | *2 hr* | *3 hr* | *4 hr* |
| **Lactate** | p=0.48 | **p<0.001** | p=0.96 | CON (*p=0.07)* | 0.36 (0.15) | 0.65 (0.19) | 0.61 (0.25) | 0.57 (0.29) | 1.00 (0.40)* | 0.88 (0.40) |
| (mmol) | | | | DOB+ † | 0.35 (0.05) | 0.64 (0.06) | 0.46 (0.12) | 0.46 (0.15) | 0.85 (0.32) | 0.75 (0.28) |
| | | | | DOB- | 0.73 (0.15) | 0.86 (0.35) | 0.72 (0.00) | 0.50 (0.10) | 1.39 (0.12) | 1.63 (0.13) |
| | | | | NE †† | 0.44 (0.13) | 0.71 (0.09) | 0.67 (0.10) | 0.63 (0.08) | 0.92 (0.31)** | 0.72 (0.20) |
| **Pyruvate** | p=0.60 | **p<0.001** | p=0.69 | CON ††† | 0.031 (0.008) | 0.023 (0.015) | 0.018 (0.013) | 0.017 (0.011) | 0.055 (0.019)*** | 0.052 (0.020)** |
| (mmol) | | | | DOB+ †† | 0.027 (0.008) | 0.031 (0.018) | 0.027 (0.017) | 0.022 (0.016) | 0.045 (0.010)* | 0.045 (0.008)* |
| | | | | DOB- | 0.038 (0.009) | 0.017 (0.006) | 0.010 (0.005) | 0.011 (0.009) | 0.080 (0.004) | 0.106 (0.010) |
| | | | | NE ††† | 0.029 (0.009) | 0.017 (0.016) | 0.015 (0.012) | 0.014 (0.010) | 0.048 (0.021)*** | 0.046 (0.018)*** |
| **Lactate/** | p=0.29 | **p<0.001** | p=0.61 | CON ††† | 11.9 (3.4) | 35.5 (15.1) | 50.0 (45.7) | 48.2 (45.6) | 19.2 (9.2)* | 16.6 (4.6)* |
| **Pyruvate** | | | | DOB+ | 13.7 (4.1) | 32.1 (29.2) | 38.4 (49.3) | 36.0 (33.6) | 19.2 (7.6) | 16.2 (3.8) |
| | | | | DOB- | 19.1 (0.9) | 59.1 (43.2) | 77.8 (35.9) | 62.1 (42.1) | 17.3 (0.6) | 15.4 (0.3) |
| | | | | NE ††† | 15.2 (3.9) | 59.5 (24.7) | 78.1 (53.4) | 75.3 (59.4) | 21.1 (7.6)* | 18.8 (5.6)* |
| **Glucose** | p=0.20 | **p<0.001** | **p=0.009** | CON ††† | 197 (86) | 92 (75) | 70 (46) | 60 (61) | 270 (146)*** | 295 (140)*** |
| (mmol) | | | | DOB+ † | 269 (182) | 120 (77) | 92 (74) | 82 (80) | 155 (55) | 175 (60) |
| | | | | DOB- | 253 (98) | 27 (3) | 91 (24) | 30 (30) | 236 (24) | 241 (41) |
| | | | | NE ††† | 187 (80) | 37 (24) | 42 (34) | 37 (27) | 170 (86)*** | 172 (88)*** |
| **Glutamate** | p=0.44 | **p<0.001** | p=0.29 | CON † | 0.0019 (0.0015) | 0.0070 (0.0031) | 0.0026 (0.0017)* | 0.0033 (0.0025) | 0.0022 (0.0024)* | 0.0015 (0.0014)* |
| (mmol) | | | | DOB+ †† | 0.0038 (0.0019) | 0.0140 (0.0111) | 0.0089 (0.0119) | 0.0070 (0.0107) | 0.0025 (0.0028)* | 0.0017 (0.0016)* |
| | | | | DOB- | 0.0017 (0.0002) | 0.0150 (0.0077) | 0.0108 (0.0025) | 0.0124 (0.0019) | 0.0026 (0.0013) | 0.0019 (0.0008) |
| | | | | NE †† | 0.0028 (0.0013) | 0.0149 (0.0123) | 0.0051 (0.0055) | 0.0027 (0.0016)** | 0.0027 (0.0016)** | 0.0021 (0.0008)*** |
| **Glycerol** | p=0.17 | **p<0.001** | p=0.35 | CON ††† | 0.012 (0.005) | 0.021 (0.003) | 0.020 (0.005) | 0.022 (0.006) | 0.052 (0.020)*** | 0.047 (0.012)*** |
| (mmol) | | | | DOB+ †† | 0.020 (0.008) | 0.023 (0.005) | 0.022 (0.007) | 0.026 (0.009) | 0.038 (0.010)** | 0.036 (0.008)** |
| | | | | DOB- | 0.022 (0.004) | 0.040 (0.020) | 0.031 (0.004) | 0.030 (0.006) | 0.060 (0.007) | 0.061 (0.007) |
| | | | | NE ††† | 0.017 (0.006) | 0.029 (0.007) | 0.030 (0.007) [a] | 0.034 (0.008) [a] | 0.053 (0.022)** | 0.057 (0.026)*** |

Values are means (SD). CON: control (*n*=8); DOB+: high-dose (i.e. ≥2.5 µg/kg/min; *n*=4) Dobutamine; DOB-: low-dose Dobutamine (i.e. ≤0.5 µg/kg/min; *n*=3); NE: Norepinephrine (*n*=7). Statistics are identical to those outlined in Table S2a. Data from DOB- were not included in statistical analyses due to only *n*=2 for microdialysis data. Between-group comparisons: [a] p<0.05 vs CON. Within-group comparison to 30 mins (i.e. pre-drug): *p<0.05; **p<0.01; ***p<0.001. Within-group effect of time from 30 mins to 4 hrs post-SCI: †p<0.05; ††p<0.01; †††p<0.001.

**Table S3.** Nonparametric data for intraparenchymal oxygenation, blood flow, cord pressure and microdialysis measures

| | GROUP | Pre-SCI *Baseline* | Post-SCI *30 min* | *1 hr* | *2 hr* | *3 hr* | *4 hr* |
|---|---|---|---|---|---|---|---|
| **Spinal cord oxygenation, blood flow and cord pressure at 1.2 cm** | | | | | | | |
| **$SCO_2$ 1.2cm** | CON | | | | | | |
| (mmHg) | DOB+ | | | 1.0 (0.7, 14.1) | 1.4 (0.7, 19.7) | | |
| | DOB- | | 0.6 (0.6, 3.3) | 0.6 (0.6, 13.2) | 0.6 (0.6, 19.3) | | |
| | NE | | | 0.6 (0.6, 1.0) | 0.6 (0.6, 0.7) | | |
| **$SCO_2$ 1.2cm** | CON | | | | | | |
| (%Δ from | DOB+ | | | | | | |
| pre-SCI) | DOB- | | | -98 (-99, -46) | -98 (-99, -22) | | |
| | NE | | | -96 (-98, -92) | -96 (-97, -56) | | |
| **SCBF 1.2cm** | CON | | | | | | |
| (a.u.) | DOB+ | | | | | | |
| | DOB- | | | 63 (56, 375) | 51 (47, 304) | | |
| | NE | | | | | 236 (177, 289) | 228 (166, 487) |
| **SCBF 1.2cm** | CON | | -64 (-74, -47) | | | | |
| (%Δ from | DOB+ | | | | | | |
| pre-SCI) | DOB- | | | | | | |
| | NE | | -78 (-84, -41) | | | | |
| **SCP 1.2cm** | CON | | | | | | |
| (mmHg) | DOB+ | | | | | | |
| | DOB- | | | 29.8 (0.4, 30.0) | | | |
| | NE | | | | | | |
| **SCP 1.2cm** | CON | | 222 (110, 457) | 226 (151, 387) | 219 (173, 357) | 219 (173, 357) | |
| (%Δ from | DOB+ | | | | | | 90 (46, 104) |
| pre-SCI) | DOB- | | | | | | |
| | NE | | | | | | |
| **Spinal cord oxygenation, blood flow and cord pressure at 3.2 cm** | | | | | | | |
| **$SCO_2$ 3.2cm** | CON | | | | | | |
| (%Δ from | DOB+ | | | | | | |
| pre-SCI) | DOB- | | | | | | |
| | NE | | | | | 38 (9, 42) | |
| **SCBF 3.2cm** | CON | | | | | | |
| (a.u.) | DOB+ | | | | | | |
| | DOB- | | | | | | |
| | NE | | | 421 (172, 3078) | 351 (171, 3315) | | |
| **SCBF 3.2cm** | CON | | | -2 (-17, 33) | | | |
| (%Δ from | DOB+ | | | | | | |
| pre-SCI) | DOB- | | | | | | |
| | NE | | | | | | |

**Table S3 *continued.*** Nonparametric data for intraparenchymal oxygenation, blood flow, cord pressure and microdialysis measures

| | GROUP | Pre-SCI *Baseline* | Post-SCI *30 min* | *1 hr* | *2 hr* | *3 hr* | *4 hr* |
|---|---|---|---|---|---|---|---|
| **SCP 3.2cm** (mmHg) | CON | | | | | | |
| | DOB+ | | | | | | |
| | DOB- | | | | | | |
| | NE | | 14.5 (7.6, 17.5) | 13.2 (8.1, 15.5) | 11.8 (7.4, 13.6) | | |
| **SCP 3.2cm** (%Δ from pre-SCI) | CON | | | | | | |
| | DOB+ | | | | | | |
| | DOB- | | | | | 16 (0, 103) | |
| | NE | | 0 (-8, 21) | -6 (-16, 29) | -13 (-18, 30) | | |
| **Microdialysis measures of spinal cord metabolism at 1.2 cm** | | | | | | | |
| **Pyruvate** (mmol) | CON | | | | | | |
| | DOB+ | | | | | | |
| | DOB- | | | | | | |
| | NE | | 0.012 (0.009, 0.015) | | | | |
| **Lactate/ Pyruvate** | CON | | | 32.0 (24.4, 51.1) | 33.3 (26.2, 50.5) | | |
| | DOB+ | | | 16.3 (11.6, 65.2) | | | |
| | DOB- | | | | | | |
| | NE | | | | | 18.9 (16.1, 23.5) | |
| **Glucose** (mmol) | CON | | | | | | |
| | DOB+ | | 151 (72, 168) | | | | |
| | DOB- | | | | | | |
| | NE | | | | | | |
| **Glutamate** (mmol) | CON | 0.0013 (0.0010, 0.0018) | | | | 0.0011 (0.0006, 0.0022) | |
| | DOB+ | | | 0.0038 (0.0021, 0.1353) | 0.0019 (0.0011, 0.0128) | | |
| | DOB- | | | | | | |
| | NE | | | 0.0028 (0.0023, 0.0196) | 0.0024 (0.0013, 0.0088) | | |
| **Glycerol** (mmol) | CON | | | | | | |
| | DOB+ | | | | | | 0.040 (0.031, 0.041) |
| | DOB- | | | | | | |
| | NE | | | | | | |

Values for non-parametric data are presented as medians and interquartile ranges (25%, 75%). Only variables with non-parametric data are included. See previous Table S2d for abbreviations. Normalcy was determined using the Shapiro-Wilk test. Grey rows or columns indicate data are not available (i.e. not collected at this time point, or insufficient *n*) for the given time point or group.
